# Harmine Selectively Drives Human Beta Cell Differentiation and Function Via Protein Kinase A Pathways

**DOI:** 10.64898/2025.12.07.692818

**Authors:** Peng Wang, Kunal Kumar, Luca Lambertini, Hongtao Liu, Olivia Wood, Shuxuan Chen, Aidan Pillard, Donald K. Scott, Joao A. Paulo, Steven P. Gygi, Susmita Khamrui, Stefan Mueller, Adolfo Garcia-Ocana, Esra Karakose, Michael B. Lazarus, Robert DeVita, Andrew F. Stewart

**Author notes:** Indicates Equal Contribution (Peng Wang, Kunal Kumar, Robert DeVita, Andrew F. Stewart). Address Correspondence to: Peng Wang PhD, Professor, Diabetes Obesity Metabolism Institute, The Icahn School of Medicine, Atran 5, PO Box 1152, One Gustave Levy Place, New York, NY 10029.

## Abstract

Harmine and other small molecule inhibitors of the kinase DYRK1A induce human beta cells to replicate and regenerate *in vitro* and *in vivo*, and are effective at reversing diabetes in animal models. In addition to its beta cell proliferative and regenerative effects, harmine also induces expression of transcription factors and other genes involved in beta cell differentiation and function, exemplified by PDX1, MAFA, NKX6.1, GLP1R, PCSK1 and many others. Harmine also rapidly enhances glucose-stimulated insulin secretion *in vitro* and *in vivo*, reversing diabetes within days in diabetic mice transplanted with a marginal mass of human islets. We had assumed that this pro-differentiation or pro-function effect was a common feature of all DYRK1A inhibitors, and was mediated by DYRK1A inhibition. Thus, DYRK1A is a primary target (we refer to it as Target 1) for harmine. Here, to our surprise, we report that the pro-differentiation effect is not a generalized action for all small molecule DYRK1A inhibitors and does not result from DYRK1A interference or genetic silencing. Instead, the prodifferentiation effect is restricted to a small select subset of DYRK1A inhibitors (harmine, 2-2c and 5-IT). Remarkably, the pro-differentiation effect results from the unique ability of these drugs to activate protein kinase A (PKA), a dual mechanism that distinguishes this class from other DYRK1A inhibitors. The beneficial effects of harmine on PKA appear to be indirect, driven by an as yet unidentified, Target 2 in the PKA pathway. These findings make it clear that all DYRK1A inhibitors are not interchangeable, and that those that drive both proliferation and differentiation/function will likely be preferable in human clinical therapeutic settings. They also provide a novel target or roadmap for enhancing human beta cell differentiation in Type 1 and Type 2 diabetes.

**Graphical Abstract:** 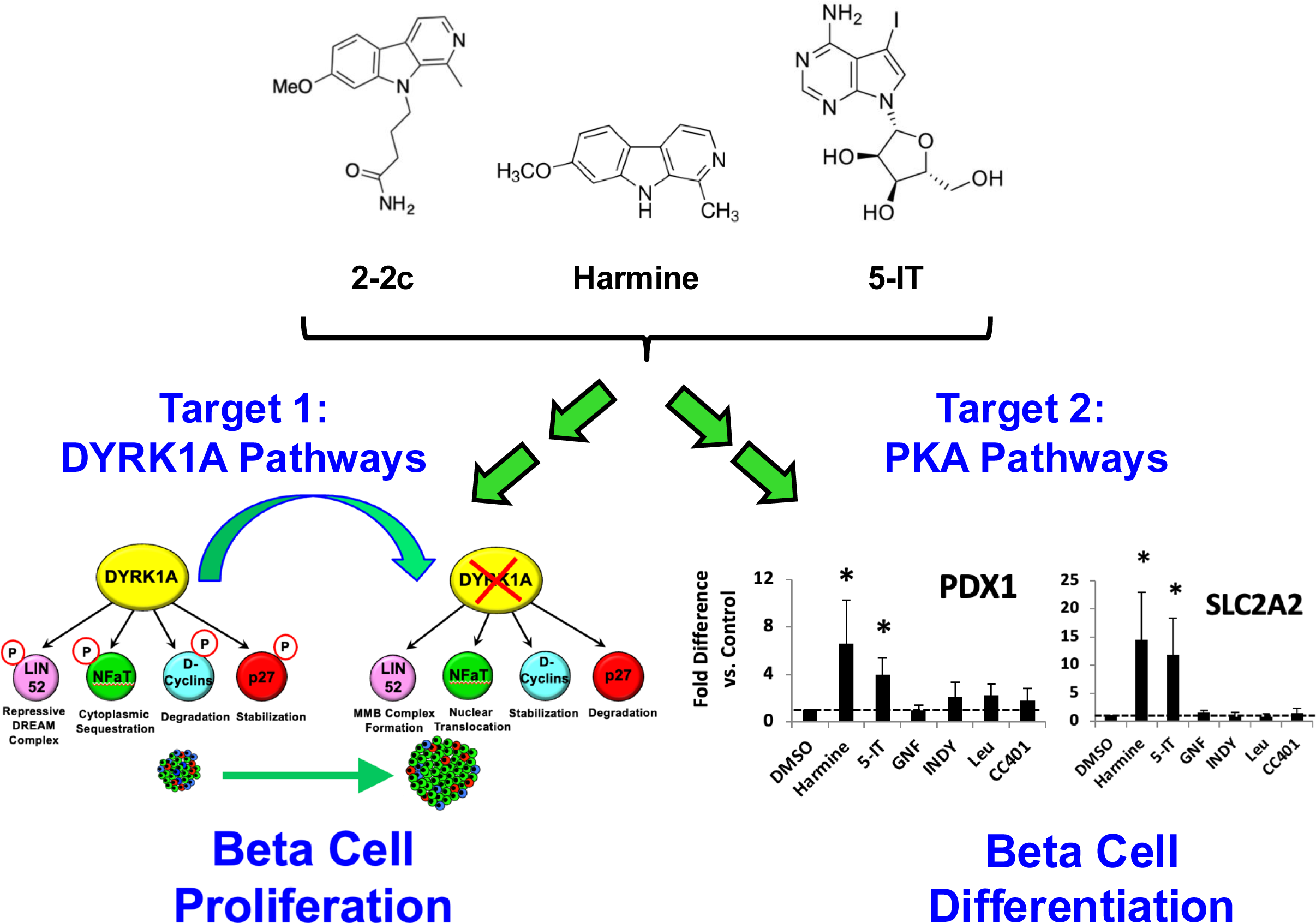

## Introduction

Diabetes affects 500 million people globally^1–6^. While insulin resistance is an underlying driver of Type 2 diabetes (T2D), both Type 1 Diabetes (T1D) and T2D ultimately result from inadequate production of insulin by the insulin-producing beta cells in the pancreas^7–11^. This beta cell anatomic and functional deficiency for all types of diabetes has prompted active programs in human beta cell replacement through whole pancreas transplant, isolated pancreatic islet transplant, and transplant of stem cell-derived human beta cells^12–16^. As an alternative, several laboratories around the world have attempted to develop drugs that are able to regenerate the beta cells that remain in most people with diabetes. Our group and others have focused on developing small molecule inhibitors of <u>D</u>ual T<u>y</u>rosine-<u>R</u>egulated <u>K</u>inase <u>1A</u> (DYRK1A) (**Fig. 1**)^17–39^. These small molecules, in *in vitro* and *in vivo* systems, are able to increase human beta cell proliferation, markedly expand human beta cell mass, increase human insulin production, normalize blood glucose and reverse diabetes in human islet transplant model systems^17,18,23^. The human beta cell proliferative effects are driven by, and require, inhibition of DYRK1A^17–20,27^, and are shared by all of the molecules shown in **Fig. 1**.

**Figure 1.**
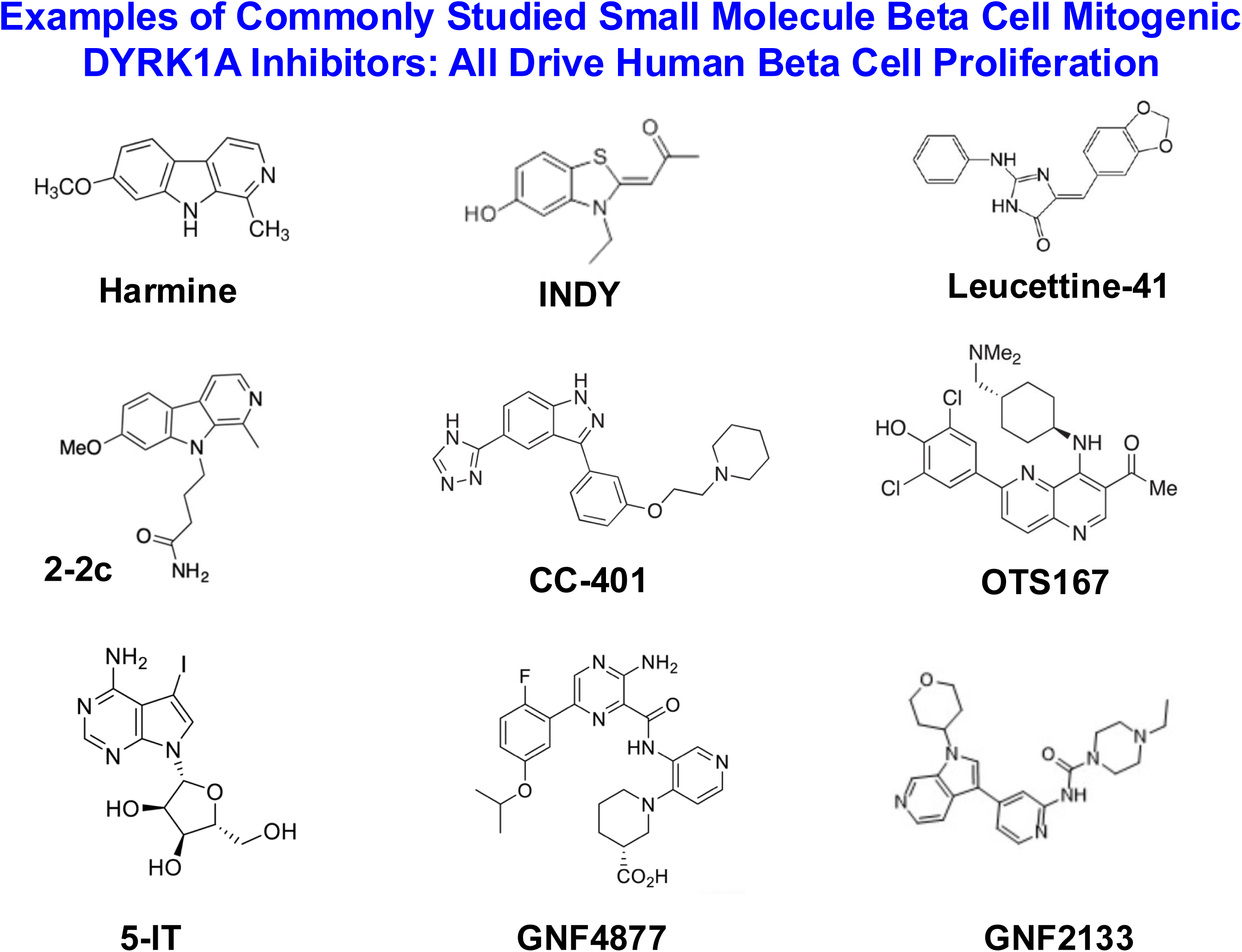
Structures of DYRK1A Inhibitors Commonly Used In Beta Cell Regeneration Research. See text for details.

An additional, remarkable, unanticipated, counterintuitive and important beneficial effect of harmine, 2-2c and 5IT is that they also enhance expression of human beta cell differentiation, manifested by increases in phenotypic or identity markers, including transcription factors (exemplified by PDX1, MAFA, NKX6.1 and many others) as well as essential beta cell functional genes (exemplified by *SLC2A2* encoding the glucose transporter GLUT2, *GLP1R* encoding the GLP1 receptor, *PCSK1* encoding the beta cell posttranslational processing enzyme for insulin, PC1/3^17,18,21,23^. These events occur at the mRNA and protein level, and, as might be anticipated, clearly enhance human beta cell function, including glucose-stimulated insulin secretion (GSIS), occur within days, long before beta cell numbers increase, and are associated with rapid (days) normalization of blood glucose in mice transplanted with an inadequate or marginal mass of human islets^17,18,21,23^. Thus, harmine, 2-2c and 5-IT (5Iodotubericidin) have two complimentary types of beneficial effects: 1) stimulation of adult human beta proliferation and expansion; and, 2) enhancing expression of genes required for human beta cell identity, function and differentiation. For simplicity, we refer to the latter collectively hereafter as “beta cell differentiation markers”.

Having dual abilities to drive both beta cell proliferation as well as differentiation makes the DYRK1A inhibitor drug class ideal for restoring the reduced beta cell mass and function characteristic of both T1D and T2D^7–11^. We had assumed initially that the proliferative and pro-differentiation effects of harmine were both attributable to interference with DYRK1A activity. In this report, we demonstrate that the pro-differentiation actions of harmine, 2-2c and 5-IT are unique among the DYRK1A inhibitor class, and do not apply to most members of the DYRK1A inhibitor class. In addition, we report that the pro-differentiation effects are entirely independent of DYRK1A, “Target 1” for all DYRK1A inhibitors. Finally, we determine that the pro-differentiation effects are due to an unknown “Target 2” of harmine, 2-2c, and 5-IT, and that this second target activates the protein kinase A (PKA) pathway, which in turn enhances human beta cell differentiation.

## Results

### Harmine, 2-2c and 5-IT Uniquely and Selectively Upregulate Human Beta Cell Differentiation Marker Genes

We have previously reported that harmine and 2-2c are able to augment PDX1, NKX6.1 and MAFA expression in human islets^18,21^, and this results in augmented human beta cell function^17,18,21,23^. Since we had assumed that this was a class effect of all DYRK1A inhibitors, we were surprised to observe that the pro-differentiation effects are limited to treatment with harmine, 2-2c and 5-IT (**Fig. 2**). In contrast, GNF4877, INDY, Leucettine-41 and CC-401, while being effective drivers of proliferation^19,27–34^, all fail to induce human beta cell differentiation markers (**Fig. 2**). Interestingly, marker induction is also selective: while the pro-differentiation effect was observed for *PDX1, MAFA, NKX6.1, SLC30A8, SLC2A2, GLP1R, SIX2, SIX3, PCSK1, ENTPD3* and *GCK*, it was not observed for other important markers of beta cell differentiation, such as the chromogranins, SLC2A1 encoding GLUT1, and *UCN3* encoding urocortin-3. **Suppl. Fig. 1** shows that the time course for induction of *PDX1, MAFA, NKX6.1, C2CD4A* and *C2CD4B* is rapid (2-4 hours), whereas *SLC2A2, ENTPD3* and *INS* only rise at later time points, consistent with their being downstream responders to *PDX1, MAFA, NKX6.1, C2CD4A* and *C2CD4B*. In contrast, *NeuroD1* and *FOXO1* showed only small and variable responses.

**Figure 2.**
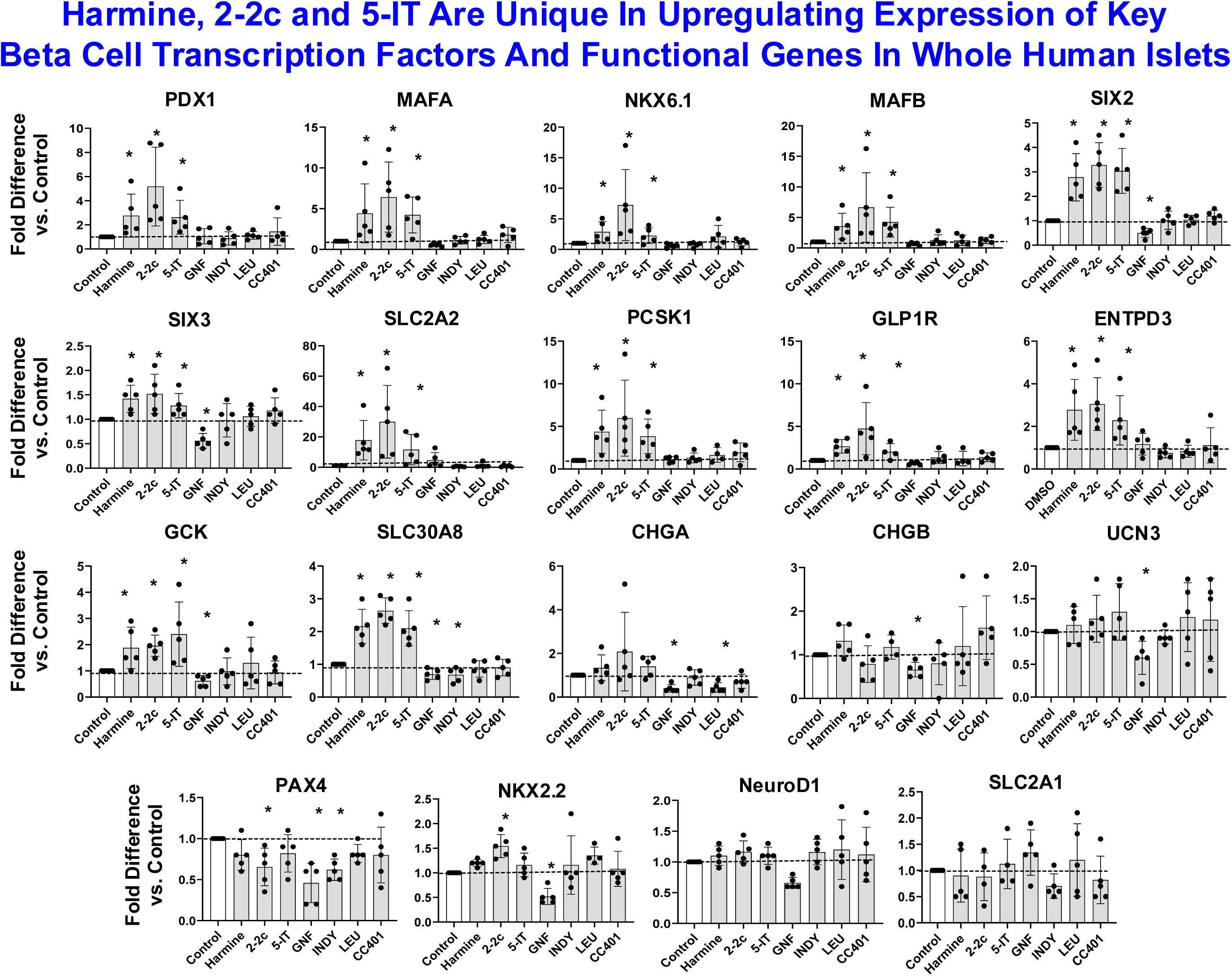
Effects of DYRK1A Inhibitors on Markers of Human Beta Cell Differentiation, Function and Identity, Assessed By Quantitative PCR on Whole Human Islets. Note that harmine, 2-2c and 5-IT reproducibly increase most of these markers in human islets, whereas GNF4877, INDY, Leucettine-41, and CC401 fail to do so. Also note that not all differentiation markers are increased by DYRK1A inhibitors, exemplified by CHGB, UCN3, PAX4, NKX2.2, NeuroD and SLC2A1 (GLUT1). Doses of each drug were selected as the most effective for human beta cell proliferation in a prior study^19^ and were: harmine 10μM, 2-2c 3μM, 5IT 1μM, GNF4877 2μM, INDY 15μM, Leucettine 41 20μM and CC-401 10μM. Asterisks indicate p<0.05 in Students unpaired T-test.

The studies In **Fig. 2** are on whole islet PCR experiments and thus include all human islet cell types. To determine if these changes were beta cell-specific, or were occurring in all islet cell types, we explored our human islet single cell RNAseq datasets^22^. As can be seen in **Fig. 3**, the increases in *PDX1, MAFA* and *SLC2A2* are almost exclusively limited to human beta cells, although there also is an increase in delta cells and cells annotated as “unknown endocrine cells”. On the other hand, changes in NKX6.1 involved several other islet endocrine cell types. This wider induction in additional islet cell subtypes also applied to other islet endocrine markers as well, such as *MAFA, MAFB, NEUROD1, C2CD4A, C2CD4B, SLC30A8*, and *NKX2.2* and *NEUROD1* (not shown).

**Figure 3.**
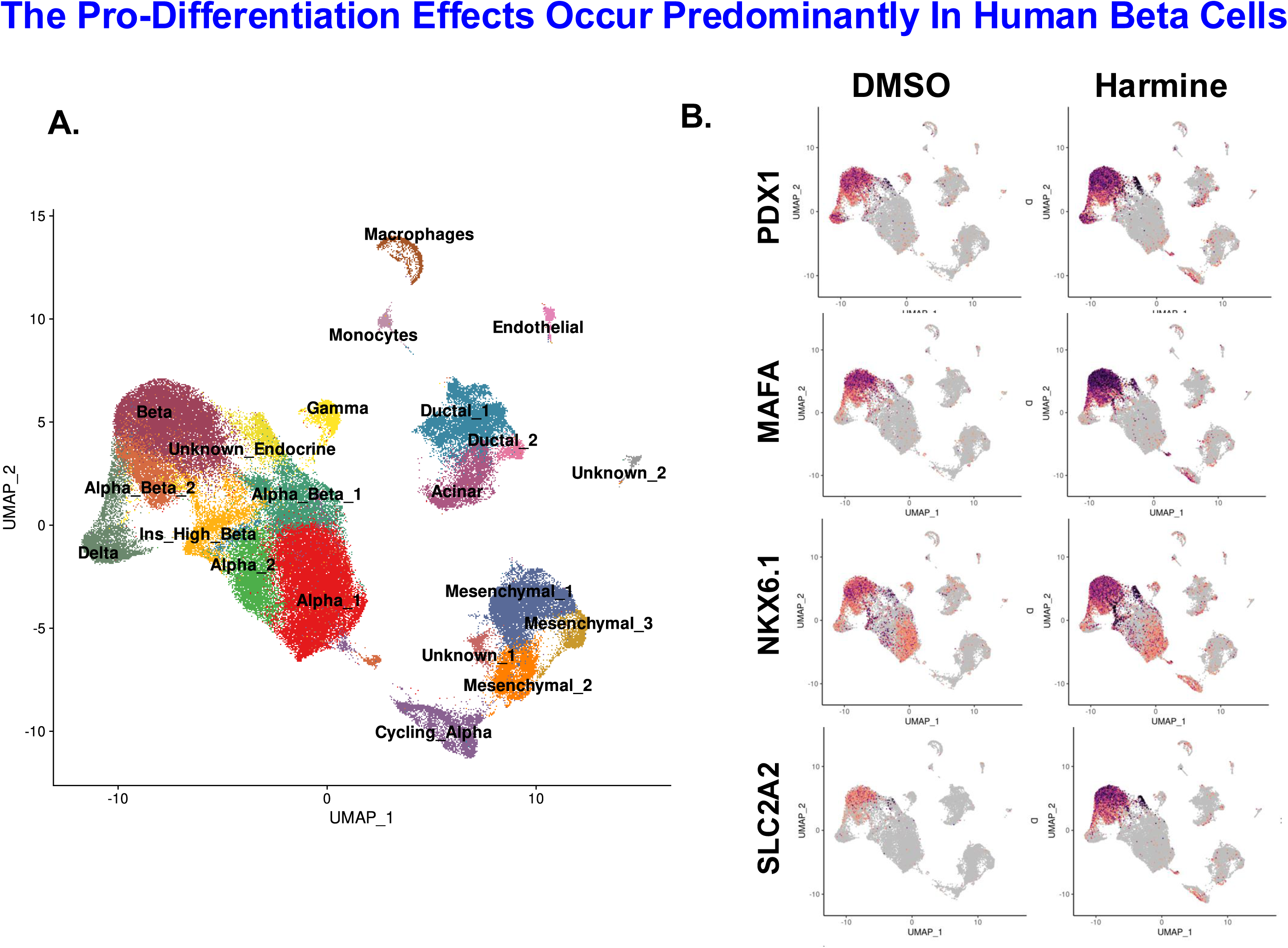
Selected Feature Plots of Single Cell RNAseq of Human Islets Treated with Vehicle or Harmine Reveal That Pro-Differentiation Events Occur Predominantly, But Not Exclusively, in Beta Cells. These feature plots are derived from an earlier study of human islets treated for 96 hours with vehicle (0.1% DMSO) or 10 μM Harmine, then subjected to scRNAseq, all as detailed^22^. N=4 human islet preps for each treatment. **Panel A** is a UMAP showing cell type annotation on a UMAP of all islet cell types. **Panel B** compares expression of PDX1, MAFA, NKX6.1 and SLC2A2 in response to vehicle vs. harmine. Note that in most cases, the major increase in gene expression occurred predominantly in beta cells, whereas in other cases, exemplified by NKX6.1, the increase was apparent in multiple islet cell types.

### Parallel Increases In Human Beta Cell Proteins

To determine whether changes at the mRNA level corresponded to the proteome, we performed proteomics on human islets treated with harmine or vehicle (0.1% DMSO). As shown in the volcano plot in **Fig. 4A**, many of the factors identified at the mRNA level translated to the proteome. For example, PDX1, NKX6.1, MAFA, C2CD4A, SLC2A2, MAFB, ENTPD3 (green circles) were all increased by harmine. Others of interest that we had not specifically queried in **Fig. 2** were also increased (SCGN, ABCC8 and HGF). In addition, markers of cell cycle activity such as Ki67, WEE1, TOP2A, CCND1 (blue circles) were also increased, perhaps as expected. Conversely, harmine treatment was associated with declines (red circles) in ALDH1A3 (a marker of beta cell stress and dysfunction^40^), CDKN1C, encoding p57^KIP2^, (a cell cycle inhibitor important in beta cell cycle arrest, and reduced with harmine treatment in prior studies^18^), and the serpin family, which has been associated with beta cell proliferation^41^. Other examples are shown in **Fig. 4A** and **Suppl. Table 3**. KEGG and GO pathway analyses did not identify additional novel pathway information. The proteomics results were independently confirmed at the immunocytochemistry level (**Fig. 4B,C**) which showed clear and statistically significant increases within beta cell nuclei of PDX1 and NKX6.1. Cell surface abundance of ENTPD3 also was increased on beta cells (**Fig. 4B,C**). These findings confirm that events observed at the mRNA level translate to the protein level, and occur in human beta cells.

**Figure 4.**
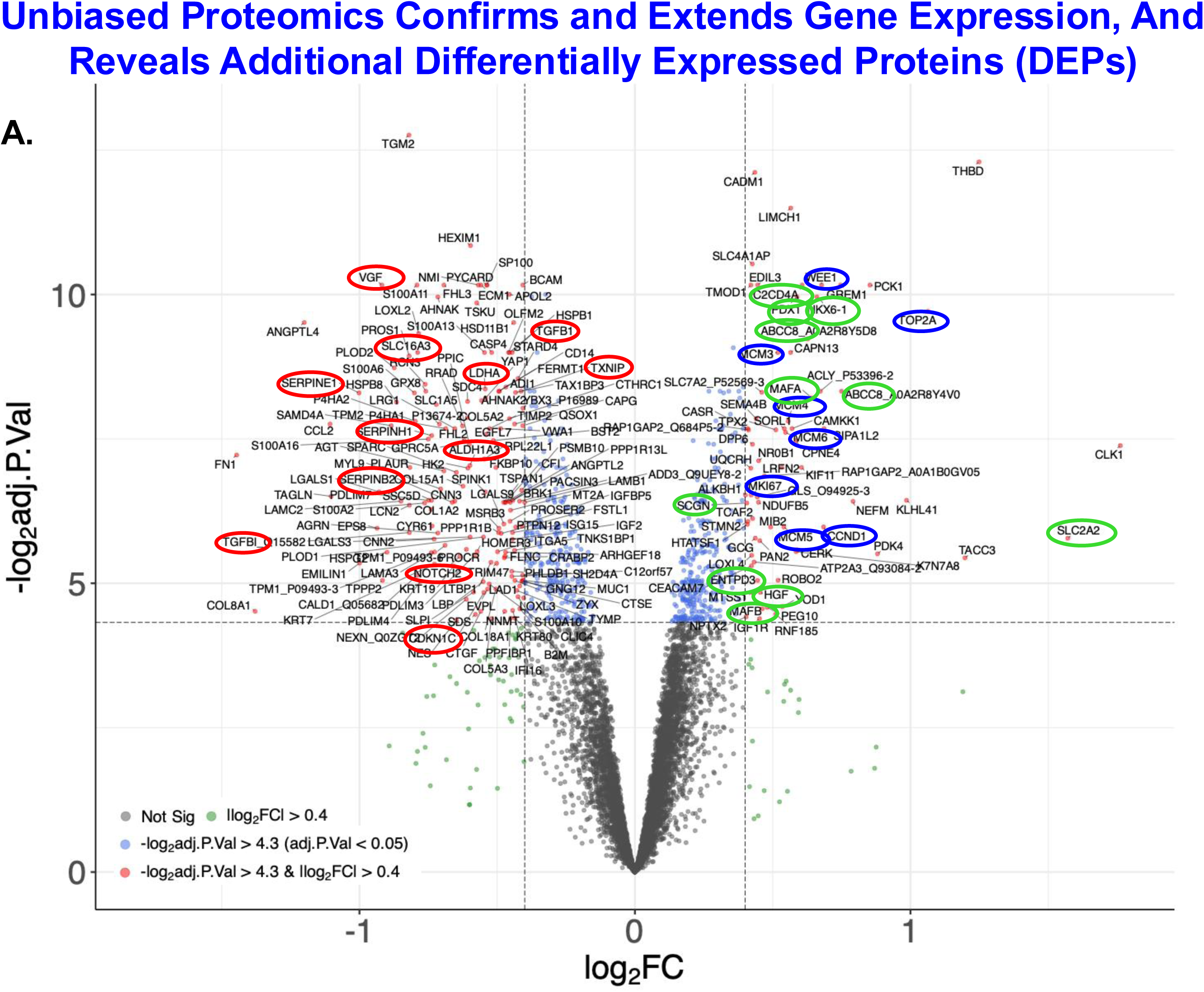
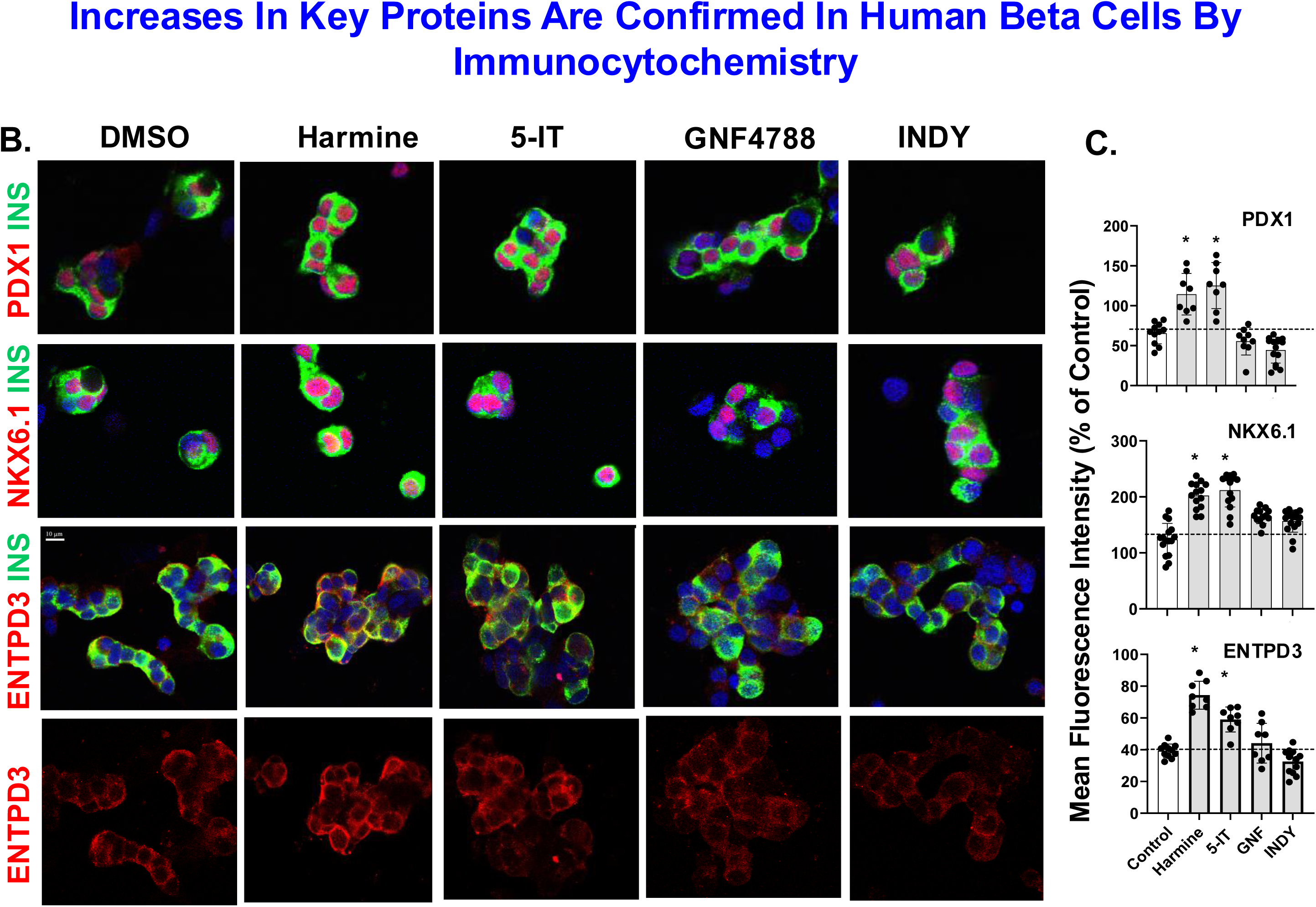
Unbiased Proteomics of Whole Human Islets and Immunocytochemistry in Human Islet Beta Cells Reveal that Changes at the mRNA Level Translate to the Protein Level. **Panal A** shows results of Harmine treatment 10μM for 72 hours on three preparations of whole human islets. Red, Blue and Green Circle are described in the text. Log-fold changes and adjusted P-values are shown on the X and Y axes. See text for details. **Panel B** shows immunohistochemistry for Insulin, PDX1, NKX6.1 and ENTPD3 in dispersed human islets. treated with the drugs indicated. Note that all three markers increase with harmine and 5-IT, but not with GNF4877 nor INDY. **Panel C** shows data from three individual experiments quantified by Image-J. Asterisks indicate P<0.05 One-Way ANOVA for repeated measures.

### The Increase In Beta Cell Differentiation Is Independent of DYRK1A

Silencing DYRK1A mimics the effects of harmine by driving human beta cell proliferation^17,19,27^. To ask whether DYRK1A inhibition also was responsible for driving differentiation, we silenced simultaneously DYRK1A and its homologue, DYRK1B, in human islets and treated the islets with control vehicle (DMSO) or with harmine. **Figs. 5A,B** demonstrate that silencing of DYRK1A and DYRK1B was effective, reducing their expression by 60%, a level more than sufficient to drive human beta cell proliferation^17,19,25,28^. In contrast, **Figs. 5C-H**, show that combined silencing DYRK1A and DYRK1B had no effect on differentiation marker expression. Critically, harmine is fully able to drive differentiation marker expression despite DYRK1A and DYRK1B silencing (**Figs. 5 C-H**). These findings make it clear that the pro-differentiation effect of harmine is not mediated by, and does not involve, DYRK1A or DYRK1B. Instead, they suggest that harmine has an additional second target that mediates its pro-differentiation effects.

**Figure 5.**
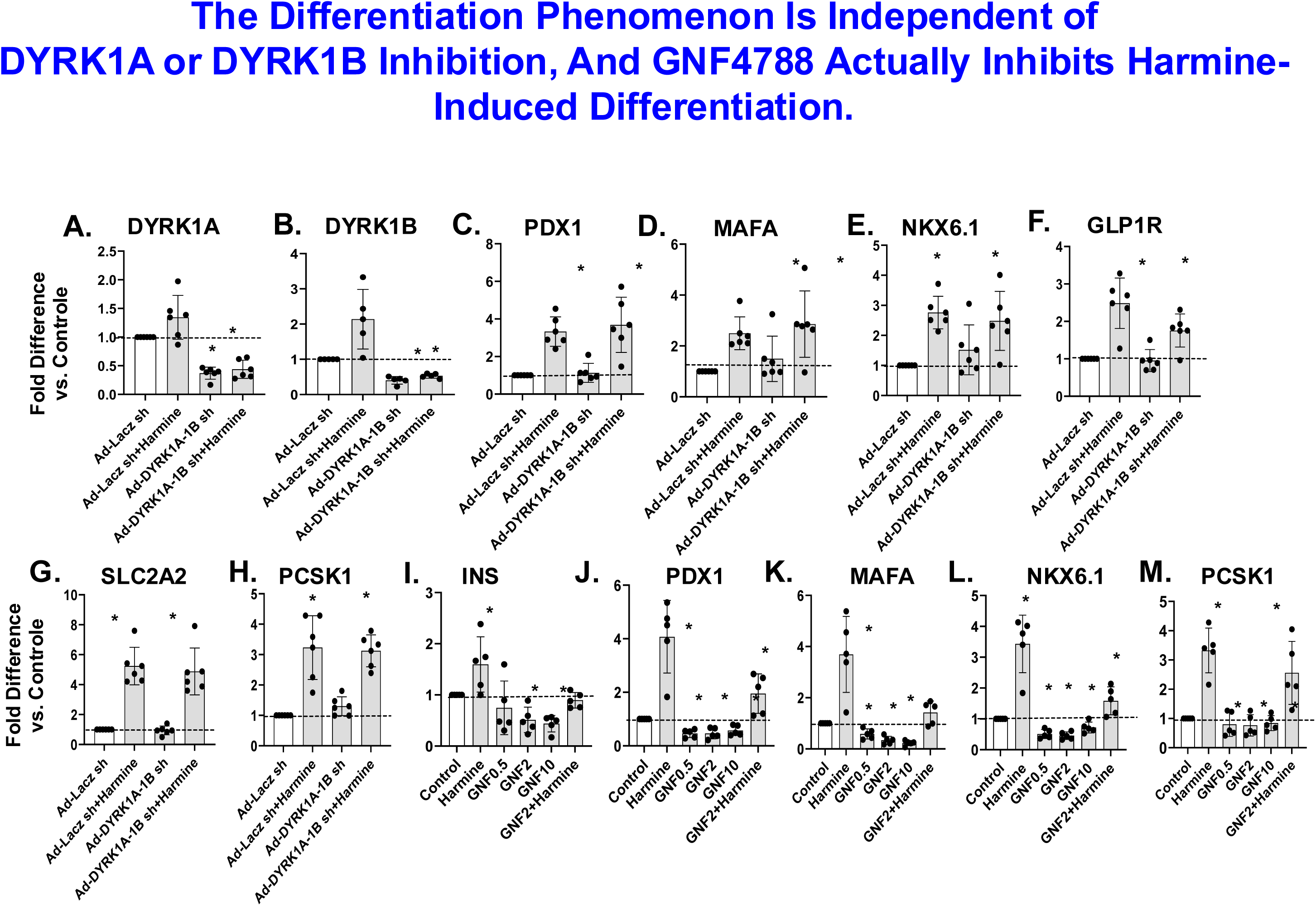
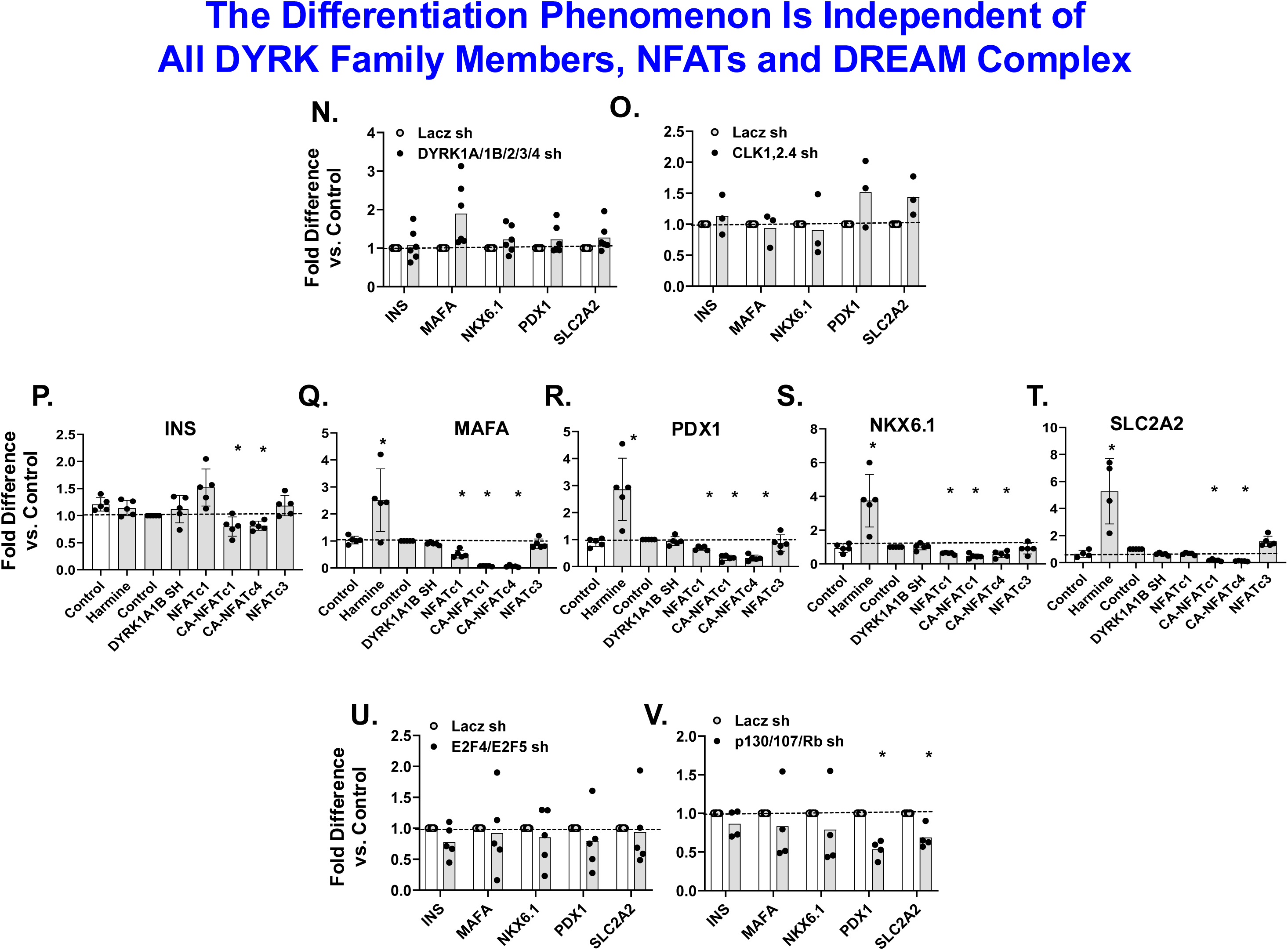
Exploring and Eliminating Candidates For The Pro-Differentiation Effects of Harmine. **Panels A,B** show that silencing both DYRK1A and DYRK1B with a single adenovirus is effective in reducing DYRK1A and DYRK1B expression by 60-70% as documented previously^17,19,25^. **Panels C-H** show that silencing DYRK1A and DYRK1B has no effect on basal differentiation marker expression. Most importantly, note that silencing DYRK1A and DYRK1B has no negative effect on harmine-induced increases in these key differentiation markers. **Panels I-M** explore the effects of GNF4877 on human islet differentiation markers. Note that while harmine displayed its usual induction of differentiation markers, GNF4877 at three different doses (0.5, 2.0 and 10μM) had no beneficial effects on differentiation markers, and in several instances actually blocked the prodifferentiation effect of harmine. **Panels N,O** demonstrate that silencing the entire DYRK family (DYRK1A, DYRK1B, DYRK2, DYRK3 and DYRK4) or CLKs 1,2 and 4 (CLK3 is not expressed in human islets) has no effect on human beta cell differentiation. **Panels P-T** show that overexpressing multiple wild-type and constitutively active members of the NFAT transcription factor family, although effective in some systems^25^, has no effect on human beta cell differentiation. Collectively, these studies make it clear that DYRK1A, DYRK1B, DYRK2, DYRK3, DYRK4, CLKs 1,2 and 4, and the NFAT family do not mediate, and are not necessary for, the pro-differentiation effects of harmine, and that GNF4877 is a nonspecific inhibitor of human beta cell differentiation. Asterisks indicate P<0.05 One-Way ANOVA for repeated measures.

### A Candidate Approach to Identifying the Pro-Differentiation Second Target

GNF4877 was one of the first DYRK1A inhibitors shown to drive human beta cell proliferation^31^. Although we and others have confirmed that GNF4877 drives human beta cell proliferation^19^, we found that GNF4877 actually inhibits both basal as well as harmine-induced human beta cell differentiation (**Fig. 5 I-M**). This is likely explained by its broad non-specificity, targeting hundreds of other kinases^19^.

Kinome-wide scans for harmine, 2-2c, 5-IT, GNF4877 and INDY indicate that they inhibit kinases in addition to DYRK1A^19^. These include DYRKs 1A,B,2, 3 and 4, and CDC-like kinases (CLKs) 1,2 and 4^19^. Thus, these become pro-differentiation candidates. We first silenced the entire DYRK1A and CLK families, and found that this had modest or no effect on differentiation markers (**Figs. 5 N,O**).

We next explored the NFAT family. Harmine fosters nuclear translocation of NFATs to the beta cell nucleus where they activate cell cycle^43,44^. We therefore overexpressed native and constitutively active forms of NFATs (c1,c3, c4) and explored effects on human beta cell differentiation markers. As with DYRK1A and CLK manipulations, these had no effect (**Fig 5 P-T**).

DYRK1A phosphorylates Ser28 on LIN52 to assemble the cell cycle repressive DREAM complex, thereby enforcing human beta cell quiescence^25^. Silencing DYRK1A, or treatment with harmine therefore induces cell cycle entry in beta cells^25^. We thus explored key members of the DREAM complex (E2Fs 4 and 5, p130 and p107) that block cell cycle entry. Remarkably, although silencing these DREAM complex members induces human beta cell proliferation^25^, it had no effect in differentiation marker gene expression (**Fig. 5U,V**). Collectively, an extensive and complete candidate exploration failed to identify a second target of harmine that might explain its pro-differentiation effects.

An Unbiased Proteomic Approach Identifies The Protein Kinase A Family as A Potential Second ProDifferentiation Target.

Having excluded known harmine-interacting candidates, we turned to an unbiased proteomic approach. We used a modified version of compound 2-2c^42^ as the bait (**Fig 6A-D**) and extracts of human islets as a proteome source. Briefly, we modified the three-carbon chain of compound 2-2c, replacing the carboxamide terminus with an amino-terminus that would allow cross-linking of the resultant compound 15 to solid phase magnetic beads. We then used this immobilized harmine analogue as “bait” in a human islet protein lysate. The bound proteins in the lysate were eluted from the solid phase with progressively increasing concentrations of 2-2c, such that the weaker binders eluted first, and higher affinity proteins eluted last. The eluted fractions were then subjected to shotgun proteomics, as displayed in **Fig. 6E**, and listed in the legend for **Fig 6**.

**Figure 6.**
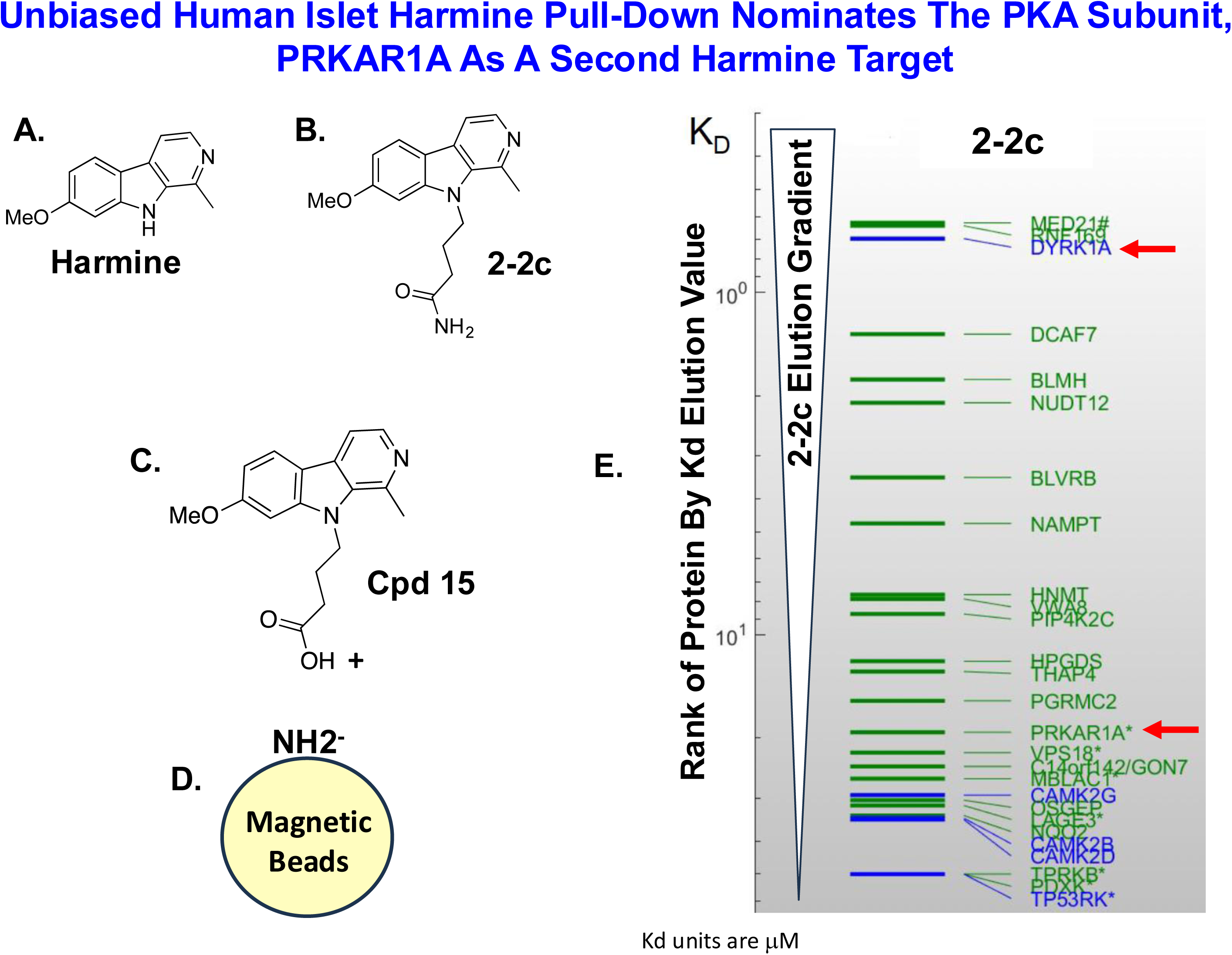
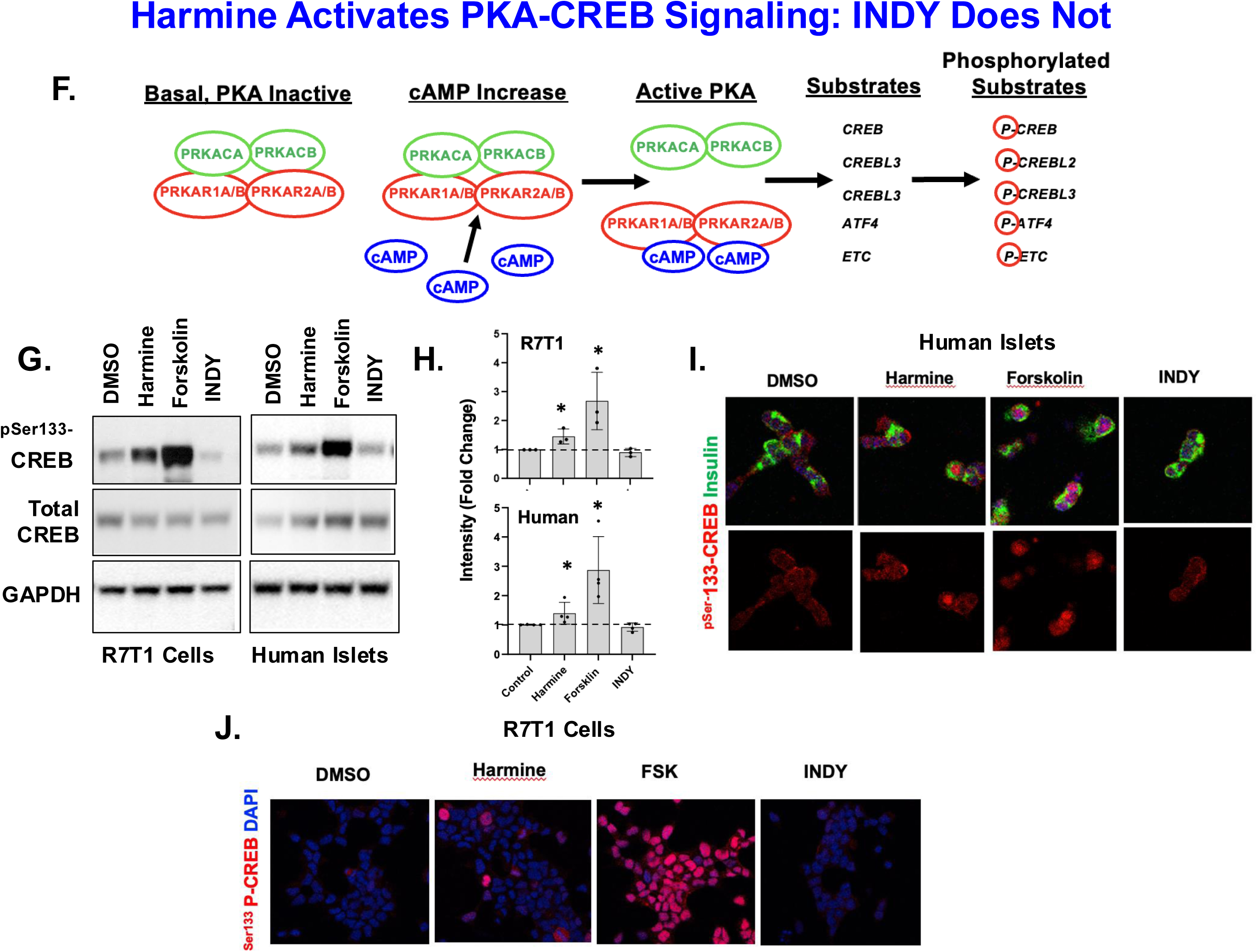
Pull-Down of Harmine-Interacting Proteins in Human Islet Extracts Reveals Protein Kinase A Interactions. **Panels A** and **B** show the structures of Harmine and 2-2c. Note that the tail of 2-2c contains a Cterminal Amide group. **Panel C**. Converting the C-terminal tail of 2-2c from an amide to a carboxylic acid in new Compound 15, permitted binding of the 2-2c/harmine-like beta carboline to solid phase magnetic beads, **D**. **Panel E.** Human islet extracts were mixed with the solid phase compound 15, and then eluted with a progressive gradient of compound 2-2c, to yield a ranked list of harmine/2-2c-binding proteins in human islets, with the strongest binders displayed at the top. The complete list, in order of affinity, higher to lowest, includes RNF169, DYRK1A, DCAF7, BLMH, NUDT12, BLVRB, NAMPT, HNMT, VWA8, PIP4K2C, HPGDS, THAP4, PGRMC2, PRKAR1A, VPS18, C14ORF142, MBLAC1, CAMK2G, OSGEP, LAGE3, NQO2, CAMK2B, CAMK2D, PRKB, PDXK, TP53RK. **Panel F.** A cartoon of the protein kinase a heterotetramer, with its association-disassociation and downstream CREB superfamily targets. **Panel G**. Harmine induces ^P133^Serine CREB phosphorylation in the R7T1 mouse beta cell line and in human islets, as does the cAMP generator Forskolin (10 μM)), whereas INDY fails to phosphorylate CREB. **Panel H.** Quantification of four experiments in human islets, with asterisks indicating P<0.05. **Panel I**. Immunocytochemistry showing that these events occur in human beta cells. **Panel J**. ^P133^Serine CREB phosphorylation Induction of ^P133^Serine CREB phosphorylation in R7T1 cells at 15 min. A complete time course is shown in **Suppl Fig. 2**.

Reassuringly DYRK1A was one of the highest affinity binders, confirming the efficacy and specificity of the approach. Other high affinity eluting proteins were a fragment Mediator 21 (MED21), RNF169 and DCAF7. The DYRK1A, DCAF7 and RNF69 signals were reproduced with multiple peptide fragments and in all triplicates. In contrast, the Mediator 21 signal was likely non-specific (only one peptide fragment in only one or three replicates). We next prepared adenoviruses to overexpress and silence RNF169, DCAF7, BLMH, NUDT12 and remaining eluting proteins in **Fig. 6E**. In none of these cases was there an effect on beta cell differentiation markers, with one exception: PRKAR1A.

**Fig. 6F** shows a model of the PKA holoenzyme complex. It is a heterotetramer composed of two catalytic subunits (PRKACA and PRKACB) and two regulatory subunits (PRKAR1A and PRKAR1B)^45^. In human islets, the most abundant members are PRKACA, PRKACB and PRKAR1A. These observations suggest that harmine may act in part by activating the PKA holoenzyme. To explore this possibility, we asked whether harmine could lead to ^133^Ser phosphorylation of the canonical PKA target protein, CREB1. Using both the mouse beta cell line, R7T1 and human islets, we found that harmine effectively leads to ^133^Ser-CREB1 phosphorylation in human islets, as does the positive control PKA activator, forskolin (**Fig. 6G,H**). Notably, the control DYRK1A inhibitor INDY, which drives human beta cell proliferation^17,19^, but not differentiation (**Fig. 2**), failed to activate PKA (**Fig. 6G,H**). These results were confirmed by immunohistochemical labeling of ^133^Ser-P-CREB1 in both human islets and R7T1 cells (**Fig. 6I,J**). **Suppl Fig 2** illustrates that the timeframe of PKA activation and CREB phosphorylation is rapid (minutes) and persists for hours. Collectively, these studies demonstrate that harmine is able to activate PKA in human and mouse beta cells, establishing PKA engagement as a shared mechanism of this distinct subgroup of DYRK1A inhibitors

### Maneuvers That Activate PKA Mimic Harmine-Induced Pro-Differentiation Effects

If harmine acts via PKA to induce differentiation, then other PKA activators should mimic the effects of harmine. To test this hypothesis, we asked whether the adenylyl cyclase activator, forskolin, could augment differentiation marker expression. **Fig. 7A** shows that this is the case: forskolin increases expression of *MAFA, PDX1, NKX6.1*, and *SLC2A2* mRNA in human islets, exactly like harmine in **Fig. 2**. **Fig. 7B-D** shows that this also occurs at the protein level in human beta cells by both immunoblot and immunocytochemistry. Importantly, INDY, does not share this prodifferentiation effect (**Fig. 7B-D**). Equally, importantly, the pro-differentiation effects of harmine in human islets can be blocked by the PKA inhibitor, H89 (**Fig. 7E,F**), providing additional documentation, along with CREB-phosphorylation, that the pro-differentiation effect of harmine is mediated by PKA.

**Figure 7.**
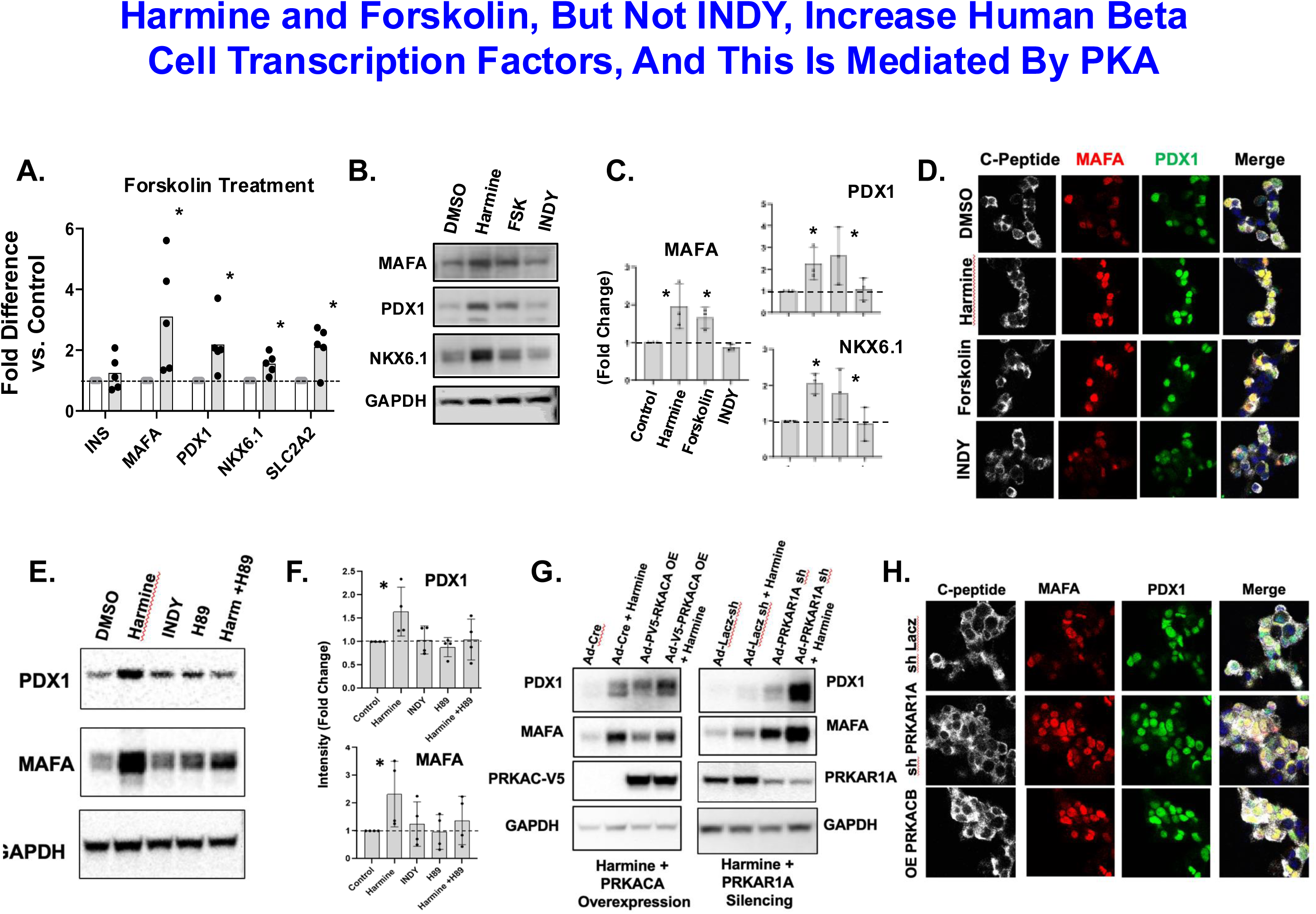
cAMP- and PKA Signaling Activates Beta Cell Differentiation Markers in Human Islets. **Panel A.** Forskolin (10μM) treatment increases expression of MAFA, PDX1, NKX6.1 and SLC2A2, mimicking the effects of harmine. **Panel B.** Immunoblots of human islet treated with vehicle, harmine, forskolin (FSK) or INDY. Harmine and forskolin, but not INDY, increase MAFA, PDX1 and NKX6.1 abundance. **Panel C.** Bar graphs of 4 experiments for Panel B. **Panel D.** Immunocytochemistry of human islets treated with vehicle, harmine, forskolin or INDY, labeled for C-Peptide, MAFA or PDX1. **Panel E.** Immunoblots of human islets treated with vehicle, harmine, INDY, the PKA inhibitor H89 (15 μM), or Harmine + H89. The key point is that harmine, but not INDY, induces PDX1 and MAFA, and this is blocked by PKA inhibition. **Panel F.** Bar graphs summarizing four different experiments similar to Panel E. **Panel G.** Left Side. Immunoblots showing that overexpression of the catalytic subunit of PKA, PRKACA, with a V5 epitope tag, increases the abundance of PDX1 and MAFA in human islets, and this is further enhanced by the addition of harmine. <u>Right Side</u>. Immunoblots showing that silencing the PKA inhibitory regulatory subunit, PRKAR1A, mimics, and is enhanced by, the effects of harmine on PDX1 and MAFA in human islets. **Panel H.** Immunocytochemistry showing that silencing PRKAR1A of overexpressing PRKACA in human islets increases expression within beta cells of PDX1 and MAFA.

### Manipulating Individual PKA Subunits Mimics the Effects of Harmine

To more deeply explore a role for the PKA subunits in **Fig. 6F**, we prepared adenoviruses to overexpress the constitutively active catalytic subunit, PRKACA, or to silence the PKA inhibitory, regulatory subunit, PRKAR1A, which we had pulled down with harmine (**Fig. 6E**). Remarkably, overexpressing the active catalytic subunit, PRKACA, led to increases in *PDX1* and MAFA (**Fig. 7G**, left panel), comparable to those induced by harmine, and this was further enhanced by harmine. More importantly, silencing the inhibitory, regulatory PKA subunit, PRKAR1A, also induced PDX1 and *MAFA* expression in human islets, similarly to harmine, and this was further enhanced by the addition of harmine (**Fig. 7G**, right panel). Finally, these events were independently confirmed in human beta cells by immunocytochemistry (**Fig. 7H**).

### The Interaction Between Harmine and PRKAR1A Is Indirect

In the conventional model of PKA in **Fig. 6F**, cAMP interacts with the inhibitory PRKAR1B subunit, resulting in dissociation of PRKAR1A from PRKACA, thereby liberating the catalytic unit to phosphorylate downstream targets such as the CREB superfamily. We reasoned that since harmine and ATP both dock in the same ATP site in DYRK1A^17,26,27^, and since the structures of cAMP, ATP and harmine share some degree of similarity, it might be possible that harmine could directly bind to PRKAR1A, in a manner similar to cAMP, thereby releasing PRKACA catalytic subunits from PRKAR1A suppression. To test this directly, we used thermal shift assays with recombinant PRKAR1A and PRKACA to determine if either was able to bind to harmine. As shown in **Fig. 8A-D**, recombinant PRKACA has a melting point of approximately 47°C, and this is not affected by harmine, indicating that harmine does not bind to PRKACA. **Fig. 8E-G** demonstrate that pure recombinant PRKAR1A has a melting point of 43°C, and this is dramatically shifted upward to 70°C in the presence of cAMP, the positive control. In marked contrast, incubation of recombinant PRKAR1A with harmine did not alter its melting point, which remained at 43°C. In aggregate (**Fig. 8H**), these findings suggest that although harmine clearly influences PKA activity, it likely does not do so through direct interaction with PRKAR1A or other members of the PKA holoenzyme.

**Figure 8.**
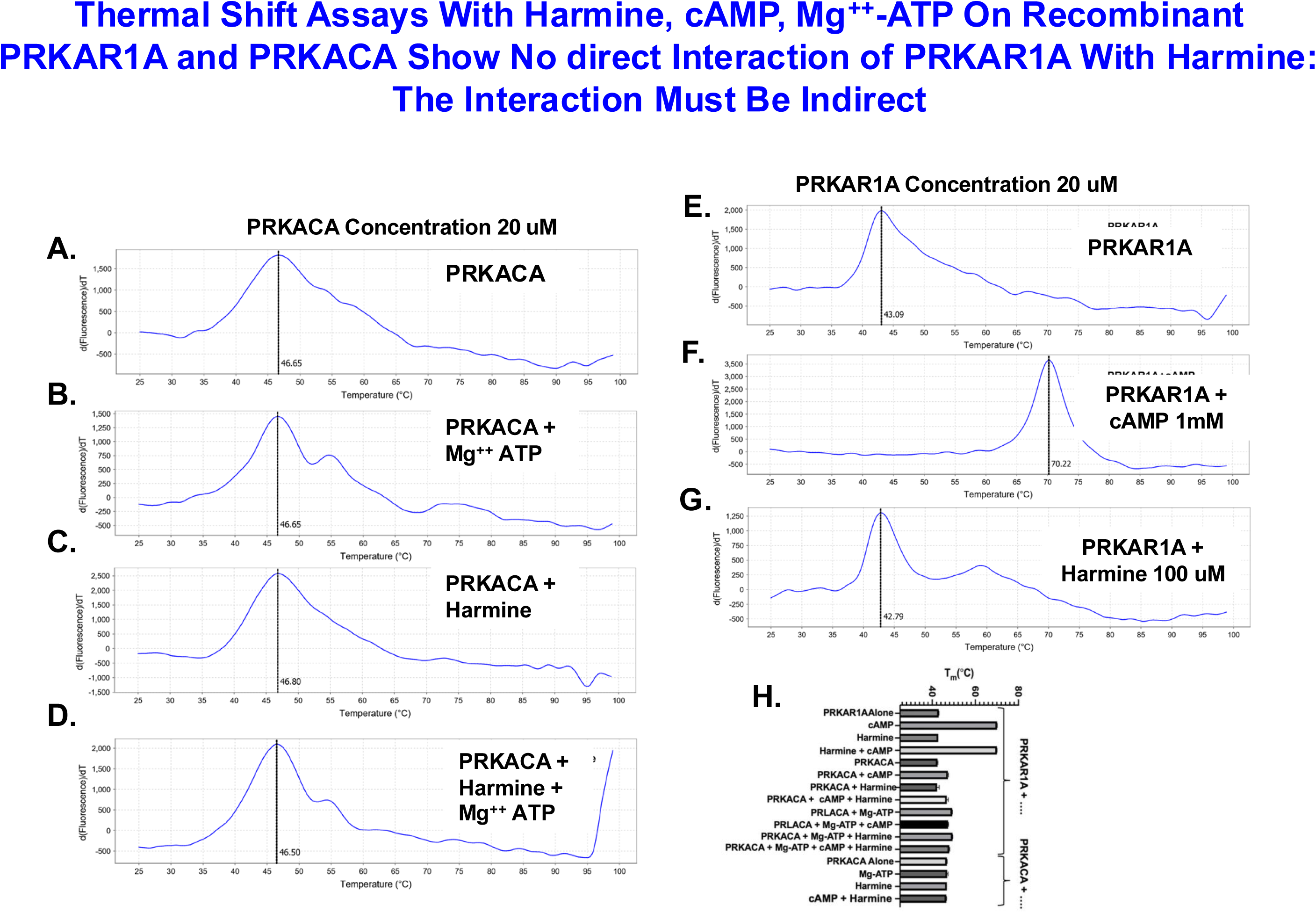
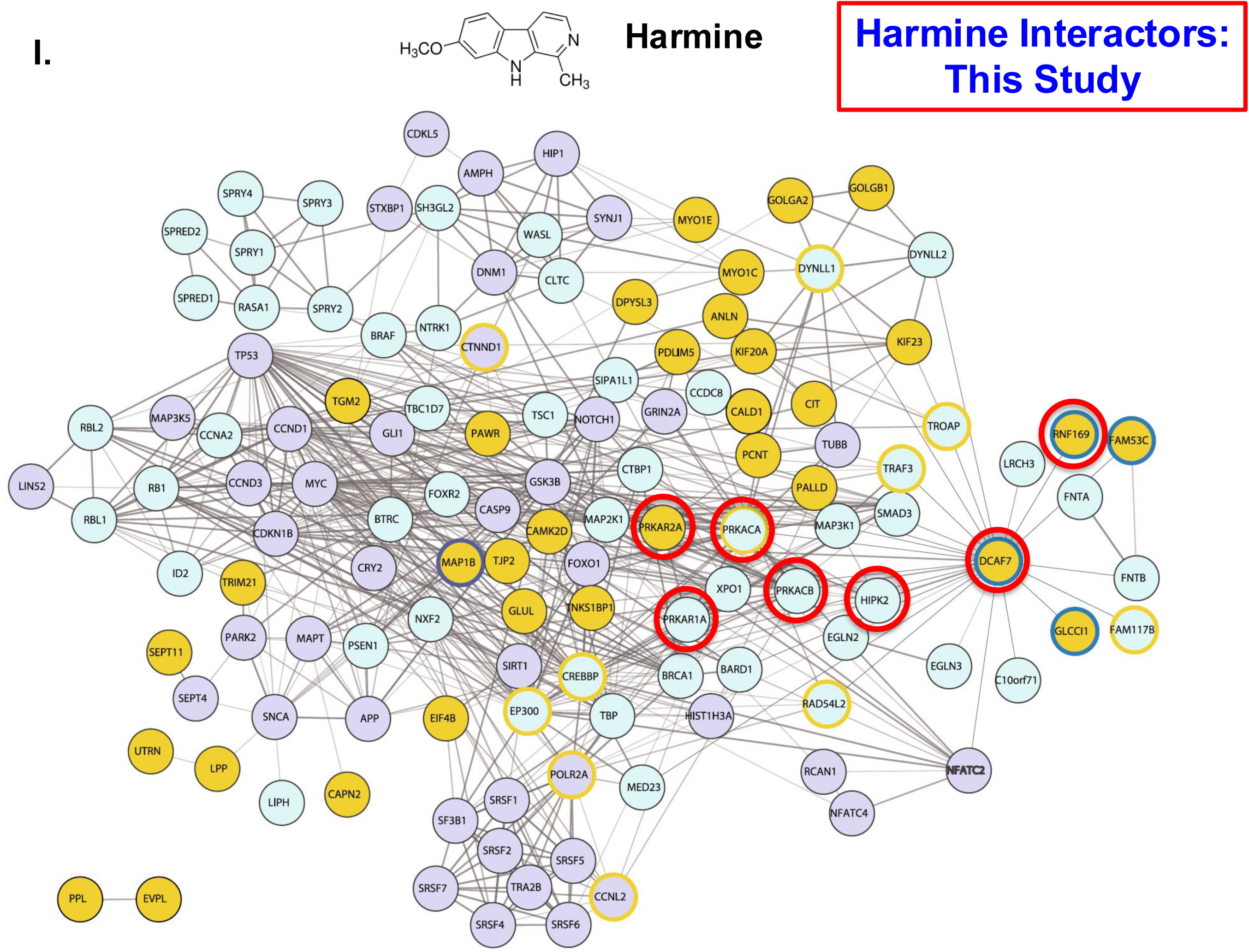
cAMP, But Not Harmine, Binds Directly to PRKAR1A, and the Harmine-DYRK1A Interactome. **Panels A-D.** Recombinant human PRKACA was assayed for thermal shift in the presence of ATP and harmine. No thermal shift was observed, suggesting that harmine, as expected, does not bind to PRKACA. **Panels E-G.** Recombinant human PRKAR1A was assayed for thermal shift in the presence of cAMP and harmine. As expected, cAMP caused a dramatic thermal shift in PRKAR1A, it’s well known target. In contrast, harmine had no effect on PRKAR1A thermal stability, suggesting there is no direct interaction between harmine and PRKAR1A, at least in vitro, and outside of the cellular context. **Panel H.** A summary of multiple thermal shift experiments. **Panel I.** An illustration of the DYRK1A interactome, and by extension, the harmine interactome, adapted from Roewenstruk and de la Luna et al^48^. The point is that many of the molecules in this cluster (DCAF7, RNF169, PRKACA, PRKACB, PRKAR1A, PRKAR2A were observed on our own pull-down data (**Fig. 6E**) and in publicly available datasets such as BioGrid and String. Note also that multiple members of the DREAM complex (LIN52, RB1, RBL1, RBL2, multiple cyclins and Myc)^25^, are present on the left side of the interactome. This may predict that harmine interacts directly with molecules such as these, and links harmine to the pro-differentiation effects in human islets.

## Discussion

These studies show that members of the DYRK1A inhibitor class of small molecules (**Fig. 1**) have two important but distinct effects on human beta cells. First, *all* DYRK1A small molecule inhibitors induce human beta cells to proliferate^17–37^. This is a central and important therapeutic property of the DYRK1A inhibitor class, can be mimicked by DYRK1A silencing^17,19,28^, and is thus a direct result of inhibition of the DYRK1A kinase. Second, a small subset of DYRK1A inhibitors (harmine, 2-2c and 5-IT) has an additional beneficial, therapeutic effect: harmine, 2-2c and 5-IT, unlike the other DYRK1A inhibitors in **Fig. 1**, also enhance human beta cell identity, differentiation and function *in vitro* and *in vivo*, defining this subgroup as a rare dual-acting class within the broader DYRK1A inhibitor family. These are therapeutically critical properties, and predict that some DYRK1A inhibitors will be more effective than others for the treatment of diabetes. Very importantly, the pro-differentiation effects of harmine, 2-2c and 5-IT are independent of DYRK1A, since silencing the entire DYRK family in human islets has no effect on human beta cell identity or differentiation status, and does not interfere with the ability of harmine to drive the pro-differentiation effect (**Fig. 5A-H,N**). Collectively, these findings demonstrate that: 1) there must be an unidentified “Target 2” for harmine, 2-2c and 5-IT - beyond DYRK1A (Target 1) - through which this group enhances human beta cell differentiation; and, 2) INDY, Leucettine and other non-differentiating DYRK1A inhibitors apparently fail to interact with this putative second “Target 2”.

We also have demonstrated that: the increases in differentiation marker gene expression occur rapidly, within 2-4 hours (**Suppl. Fig 1**); they are confined principally to beta cells in single cell RNAseq datasets (**Fig. 3**); the increases at the mRNA level translate to the protein level, by unbiased proteomics, by immunoblot, and by immunocytochemistry (**Figs. 2-4**); and, these changes were not observed with the non-differentiating DYRK1A inhibitor, INDY (**Figs 2,6,7**). This suggests that INDY may not engage Target 2.

Searching for the novel “second target”, we began with an extensive candidate approach, eliminating previously reported candidates and pathway such as DYRK1A, DYRK1B, DYRK2, DYRK3, DYRK4, the DREAM complex (p107, p130, E2F4), CLKs1,2 and 4, and the NFAT family (**Fig. 5**), all reviewed previously^17–39^. We also demonstrated that one particularly promiscuous, non-selective DYRK1A inhibitor, GNF4877^31^, actually blocked harmine-induced differentiation, likely reflecting its promiscuous interference with multiple important kinases (**Fig. 5I-M**).

Since the candidate approach was unrevealing, we next employed an unbiased pull-down proteomic approach hoping to identify a “Target 2” in lysates of human islets, using a tethered analogue of harmine/2-2c as bait^27^. This identified 24 proteins, including DYRK1A (an important positive control), as well as multiple novel harmine-interacting proteins, including DCAF7, RNF169 and many others (**Fig. 6E**). We overexpressed and silenced these proteins and assessed effects on differentiation in human islets, but found none, until we reached PRKAR1A. To our surprise, harmine (but not INDY) mimicked canonical PKA effects on ^133^Ser-CREB phosphorylation in human islets and beta cells, and the mouse beta cell line, R7T1 (**Fig. 6G-J**). Conversely, cAMP/adenylyl cyclase activation with forskolin mimicked the beneficial effects of harmine on human beta cell differentiation, and these effects of harmine were blocked by the PKA inhibitor, H89 (**Fig. 7A-F**). Finally silencing the PKA inhibitory subunit that we had pulled down in **Fig. 6E**, PRKAR1A, activated differentiation in human islets and beta cells, as did overexpressing the catalytic subunit, PRKACA (**Fig. 7G,H**). Together, these findings make a convincing argument that harmine recruits and activates PKA signaling via physical interaction with PRKAR1A, and that PRKAR1A might be “Target 2” for harmine.

We were surprised to observe, using thermal shift assays on pure recombinant PRKAR1A or PRKACA, that there is no apparent direct interaction between harmine and PRKAR1A (**Fig. 8A-H**). This may suggest that PRKAR1A is not the principal “Target 2” for harmine, but that harmine may act indirectly on another unknown “Target 2” protein that modulates PKA holoenzyme activity. It is also possible that harmine and PRKAR1A may interact in live cells in their native context, but that other cellular proteins or co-factors required in cellular context are lost in cell lysates, or with recombinant proteins *in vitro*. Exploring these and other possibilities will require extensive additional studies. Alternately, or in addition, harmine may act indirectly through another component-protein or protein complex - that modulates PKA holoenzyme activity. Importantly, the data also do not exclude a mechanistic alternative in which harmine, 2-2c ad 5-IT induce conformational or steric changes in DYRK1A itself, leading to altered assembly of the DYRK1A-associated interactome. In this scenario, the biological effects could arise from changes in complex composition unique to this subgroup, even though all DYRK1A inhibitors bind the same ATP-site catalytic target.

In line with a greater DYRK1A / harmine interactome concept, in the pull-down experiment with harmine (**Fig. 6E**), we identified DCAF7 and RNF169 as the highest affinity binders of harmine. Review of proteomics web tools like BioGrid^46^, and String^47^ reveal that DYRK1A, DCAF7 and RNF169 are frequent interactors in multiple cell types. As one example, Roewenstrunk and de la Luna^48^ identified a broad network of DYRK1A interactors in HeLa cells, shown in modified form in **Fig. 8I**. As another example, Litovchick et al have observed similar findings^49^. As yet another example, Li and Mohan have shown that DYRK1A interacts with EP300 and CBP/CREBBP to modulate H3K27Ac abundance at enhancers^50^. Collectively, these reports illustrate the fact that there is a large potential “harmine interactome” of protein partners in human beta cells through which harmine might interact to activate PKA signaling and to drive human beta cell differentiation. This interactome may include RNF169, DCAF7, DYRK1A, the CREB/CBP superfamily, FAM53C, FAM117, multiple PKA superfamily members (PRKAR1A, PRKAR2A, PRKACA, PRKACB) and many others as reflected in **Fig. 8I**. Extensive additional studies will be required to elucidate which, if any, of the proteins is a direct target of harmine, 2-2c and 5-IT. Finally, it is important to emphasize that while DYRK1A is a recurrent member of the DYRK1A interactome^46–50^, DYRK1A *per se* not required for the pro-differentiation effect, since silencing DYRK1A and other DYRK family members does not interfere with the pro-differentiation effects of harmine (**Fig. 5A-H,N**).

It was entirely unexpected to learn that harmine can activate PKA signaling. GLP1RAs, which act in part through PKA pathways, do not activate adult human beta cell proliferation^51^, though they are able to activate juvenile human beta cell proliferation^51^. While limited to rodent cell lines and animal models, studies by Hussain et al^52^, Wicksteed et al^53^, Seino et al^54^ and Li et al^55^, all indicate that activating the cAMP-PRKAR1A-CREB-CBP pathway can induce mouse beta cell proliferation or release of insulin secretory granules.

GLP1RAs have little or no effect on human beta cell differentiation^18^. Thus, it was a surprise to observe that harmine (or any other DYRK1A inhibitor) might induce human beta cell differentiation programs via activation of PKA signaling. In support of a harmine-PKA-differentiation axis, Montminy et al, in rat Ins1 beta cells, suggested that CREB can recruit the CREB coactivator, EP300, to promoters or enhancers of the islet/beta cell pioneer transcription factor, NeuroD1^56^. Minikawa et al suggested that PKA can enhance mouse beta cell differentiation by altering histone (H3K9) methylation^57^. Aida et al have suggested that PKA signaling is required for MAFA induction in mouse beta cells^58^. In a rare human islet study, Tiemann et al have suggested that cAMP-PKACREB signaling serves to repress the alpha cell transcription factor, ARX1^59^. In contrast to the studies above, there are no prior reports of PKA-induced differentiation of human beta cell lineage or differentiation markers.

PKA is a complex molecule, as are its multiple subunits. In one model, Lu et al recently described a 3-D structure of the PKA heterotetramer^45^, challenging the conventional concept that PKA subunits associate and disassociate, suggesting instead that changes in tertiary structure of the holoenzyme, driven by cAMP, result not in subunit dissociation, but rather reconfiguration of the entire heterotetramer in a manner that fosters its catalytic activity. This effect would be overlooked in studies using isolated recombinant single proteins, as in our thermal shift assays. As a second example of PKA complexity, Lee et al^60^ and Hardy et al^61^ have suggested that PRKAR1A exists individually in subcellular liquid-liquid phase separated (LLPS) bodies in various intracellular compartments that may serve as local cAMP reservoirs and/or buffers. As a third example of PKA complexity, PKA exists in multiple subcellular domains and forms: cell membrane-associated, mitochondria-associated, nuclear/nucleolar, and ER/Golgi associated, etc. This subcellular localization and trafficking is governed in part by the AKAP (A-Kinase Anchoring Protein) family of PKA chaperones^62^. Where specifically, in a subcellular sense, harmine might be affecting PKA activity to enhance beta cell differentiation is unknown. Moreover, which are the relevant CREB/ATF superfamily members, which co-activators (e.g., P-300, CBP, CRTC) might be required, which specific genes and/or enhancers they target, and, how these might activate differentiation markers, all remain unknown? These intriguing questions will provide fruitful areas for future research.

This study leaves important questions unanswered. First, what is the hypothetical “Target 2” for harmine? Is it a protein or other class of molecule? Could it represent a discrete protein or other regulatory component, or might the observed effects instead arise because this subgroup of DYRK1A inhibitors induces conformational or steric changes in DYRK1A that alter the DYRK1A-associated interactome? Such changes in complex assembly could recruit different partners into the DYRK1A signaling network and might explain why harmine, 2-2c and 5IT activate pathways that other DYRK1A inhibitors do not, even though all bind the same catalytic target. Does it have a different K_d_ for harmine, 2-2c, 5-IT vs. other DYRK1A inhibitors? Or, might it really be PRKAR1A, but its physiological interactions require intact cell context? Second, how does “Target 2” - or alternatively, these DYRK1A-dependent complex-assembly effects - mediate pro-differentiating effects of harmine? Might “Target 2” be a useful drug target for novel beta cell regenerative/functional strategies? Could components of this process represent useful entry points for novel beta-cell regenerative or functional strategies? Third, how specifically does harmine interact with the greater harmine-DYRK1A interactome in **Fig. 8I**, and which nodes in this network are selectively influenced by this subgroup of compounds? Fourth, the PKA structural model in **Fig. 6F** will continue to evolve over time. As it does, will its interactions with the harmine-DYRK1A interactome will become clearer? And which specific interactome members are required, and how do they fit into the pro-differentiation story? Fifth, what are the roles of DCAF7, RNF169, and the other recurring DYRK1A, harmine-interacting proteins in **Fig. 8I**? Many have been reported to exist in the “DYRK1A interactome” on other cell types, but their biology in human beta cells is entirely unexplored.

In conclusion, we report the surprising observation that the adult human beta cell mitogenic and proliferative molecules, harmine, 2-2c and 5-IT, selectively and uniquely enhance human beta cell function. While the pro*<u>proliferative</u>* effects of harmine and all DYRK1A inhibitors is attributable to a combination of effects on DREAM complex, NFATs, cyclins and CDKs, the selective induction of human beta cell *<u>differentiation</u>* by harmine, 2-2c and 5-IT has not been reported previously. These findings have clear and very important therapeutic implications: the optimal human beta cell therapeutic agent will be a DYRK1A inhibitor that drives *both* proliferation *and* differentiation. In addition, while it has long been apparent that cAMP/PKA signaling may drive beta cell proliferation, it has never been suggested that harmine may directly or indirectly interact and activate any member of the larger PKA pathway, or that harmine may employ PKA signaling to enhance cell differentiation. Going forward, the precise biochemical identity of the direct “Target 2” of harmine, and the roles of other members of the “DYRK1A interactome” in human beta cells need to be identified, as do the specific downstream mediators of PKA that mediate the pro-differentiation effects might be (which CREBs, ATFs, etc). These may also lead to an even better next generation of human beta cell regenerative drugs. These are exciting questions for future study.

## Acknowledgments

The authors wish to thank Bonnie and Joel Bergstein, Lonnie and Thomas Schwartz, and Martha and Fred Farkouh families for their constant support of this research. We thank Stefan Mueller at Evotec in Munich, Germany, for help with proteomics. We also thank the NIDDK-supported Human Islet and Adenovirus Core (HIAC) of the Einstein-Sinai Diabetes Research Center (ES-DRC), and the NIDDK Integrated Islet Distribution Program (IIDP), Prodo Laboratories, Patrick MacDonald at the Alberta Diabetes Institute Islet Core at the University of Alberta in Edmonton (www.bcell.org/adi-isletcore) with the assistance of the Human Organ Procurement and Exchange (HOPE) program, Trillium Gift of Life Network (TGLN), and other Canadian organ procurement organizations. We also thank Dr. Tatsuya Kin at the University of Alberta, Edmonton, Alberta, Canada, Dr. Piotr Witkowski at the University of Chicago, Chicago, Illinois, and Dr. Fouad Kandeel at the City of Hope Medical Center, Duarte, California, for providing human organ donor islets. This work was supported by NIH grants P30DK020541, R01DK116873, R01DK116904, R01DK125285, R01DK105015, R01DK129196, R01 GM132129, R01 DK139631, R35GM124838, R35 CA232128, and P01 CA203655.

## Author Contributions

P.W., K.K., R.J.D. and A.F.S. conceived of the studies. P.W., O.W., H.L., A.P., S.C., M.B.L., S.K., E.K., performed experiments. P.W., L.L., A.F.S. analyzed data. P.W., and A.F.S. wrote the manuscript. K.K., R.J.D., D.K.S. provided reagents and advice. A.G.O., E.K., reviewed data and provided advice.

## Guarantors

P. W. and A.F.S. guarantee the data in this manuscript

## Materials and Methods

### Human Islets

Isolated, de-identified human pancreatic islets from otherwise normal organ donors were provided by the NIH Integrated Islet Distribution Program (IIDP, https://iidp.coh.org), Prodo Laboratories, The Alberta Diabetes Institute, and the Transplant Surgery Department, University of Chicago. Details and demographics of the donors and islet preparations are provided in **Suppl. Table 1**. In all, a total of 85 human islet preparations were studied. Donor ages ranged from 16 to 71 years old. The mean age (±SEM) was 45.9 ± 13.6 years; mean BMI (±SEM) was 28.8 ± 5.5 kg/m^2^ (range 10-47.6). 62 donors were male and 23 were female; 48 were White, 10 Hispanic, 5 Asian, 12 Black, and race was not reported for 9. Mean cold ischemia time was 469.8 ± 182.2 minutes (range 76-1020 minutes, excluding “not provided”). HbA1c values ranged from 4.2% to 6.3%, with a mean of approximately 5.4%. Islet purity ranged from 75% to 99% (mean ± SEM 86.04% ± 5.68%). Causes of death included stroke, head trauma, anoxia, and cerebrovascular accident (CVA).

### Chemicals

Accutase (Mediatech: 25-058-CL), harmine (286044, Sigma), leucettine-41 (MR-C0023, Adipogen), INDY (4997, Tocris Biosciences), 5-Iodotubercidin (5-IT) (1745, Tocris), Recombinant human TNF-α (210-TA010, R&D Systems), GNF4877 (HY-129492, Medchem Express), CC-401 (HY-13022A, Medchem Express), Forskolin (Selleckchem), H89 (Medchem Express). 2-2c was synthesized in The Drug Discovery Institute and The Department of Pharmacological Sciences, The Icahn School of Medicine at Mount Sinai.

### Adenoviruses and Transduction

For adenoviral silencing, four target shRNA sequences for each target gene were designed using the Thermo Fisher RNAidesigner online tool. Target sequences were cloned into Block-iT U6 RNAi entry vector (K494500, Invitrogen), and silencing efficiency was evaluated in HEK293 cells. The most effective target sequences were used to generate Ad-shRNAs in the Block-iT adenoviral RNAi vector (K494100, Invitrogen). The virus that silences both DYRK1A and DYRK1B simultaneously has been reported previously^17,19,21^. Adenoviruses for silencing RBL2/p130/RBL1/P107/Rb1, E2F4/ E2F5, and CLK1/CLK2/CLK4 have been reported previously^25^. Ad.NFATC1 has been previously described^25^. Mouse Ad.ca-nfatc1 and ca-nfatc2 were gifts from Drs. Alan Attie and Mark Keller, Department of Biochemistry, University of Wisconsin. NFAT4 (NFATC3), CA-NFAT2 (CA-NFATC1), CA-NFAT3 (CA-NFATC4) are described in detail in reference^25^. AdshRNA-PRKAR1A was prepared using U6 RNAi entry vector (45-0511, Invitrogen). The PRKAR1A shRNA target sequences is GGGATAACTTCTATGTGATTG. The Ad-V5-PRKACA construct was generated by PCR amplifying PRKACA from the Addgene pDONR223-PRKACA plasmid (Cat. #82311), using a forward primer containing an N-terminal V5 epitope tag. The tagged PCR product was cloned into a Gateway destination vector under the control of the RIP promoter.

### Human Islet Dispersion and Virus Infection

Islets were centrifuged at 1500 rpm for 5 min, washed twice in phosphate-buffered saline (PBS), re-suspended in 1 ml of Accutase and incubated for 10 min at 37°C. During digestion, the islets were dispersed by gentle pipetting up and down every 5 min for 10 sec. Complete RPMI medium containing 11 mmol/L glucose, 1% penicillin/streptomycin with 10% fetal bovine serum (FBS) was then added to stop the digestion. Dispersed cells were then centrifuged for 5 min at 1500rpm, the supernatants removed, the pellets re-suspended in complete medium, and the cells then plated on coverslips with 30 µl of cell suspension per coverslip. Poly-D-Lysine/Laminin-treated cover slides or chamber slides were used. Cells were allowed to attach for 2 hr at 37°C or were transduced with adenovirus for 2 hr. After 2 hr, 500 µl complete RPMI medium was added in each well to terminate adenoviral transduction. Cells were cultured for 24-96 hr as described in the Figure Legends. Detailed protocols are provided in reference^63^.

### Compound Treatments

For compound treatments, dispersed islet cells were allowed to recover on coverslips for 24 hr. RPMI complete medium was then replaced with fresh medium containing harmine or vehicle (0.1% DMSO) for 72-96 hr. Doses used were derived from publications that demonstrated proliferation in human beta cells^17,19, 25,27,28,31,33,34^, and were harmine (10 μM), 5-IT (1 μM), GNF4877 (2 μM), INDY (15 μM), Leucettine-41 (20 μM), CC-401 (10 μM), Forskolin (10 μM) and H89 (15 μM). For the combination treatment with adenovirus and compounds, islet cells were transduced first, and then 24 hours later the cells were treated with compound.

### qPCR Methods

RNA was isolated and quantitative RT-PCR was performed as described previously (3). Gene expression in dispersed islets was analyzed by real-time QuantStudio Real-Time PCR System. Primer sequences are listed in **Suppl. Table 2**.

### Immunoblotting Procedures

Immunoblots were performed on whole human islets as described in detail previously^17–20,25^. Primary antisera were: PDX1 (Ab219207, Abcam), MAFA (Ab264418, Abcam), NKX6.1 (DSHB, F55A10-c), V5 (46-0705, Invitrogen), PRKAR1A (MA5-24981, Invitrogen), CREB (67927-1-Ig, Proteintech), pSer133-CREB (9198s, Cell Signaling), and GAPDH (2118s, Cell Signaling).

### Immunoblot Quantification

Western blot bands were quantified using ImageJ (NIH). Blot images were converted to 8-bit and analyzed without altering contrast. Identical rectangular areas were drawn around each band, and integrated density was measured. Background intensity was taken from a nearby region and subtracted from each band. Target protein signals were normalized to their corresponding loading control and then expressed relative to the control condition, which was set to 1.0.

### Immunocytochemistry and Antisera

Immunocytochemistry was performed on coverslips fixed in fresh 4% paraformaldehyde. Accutase-dispersed human islet cells were plated on coverslips as previously described (35). Primary antisera were: C-peptide (DSHB GN-ID4); PDX1 (Ab47308, Abcam), NKX6.1 (DSHB, F55A10-c), ENTPD3 (provided by Dr. Alvin Powers, Vanderbilt University), and pSer133-CREB(9198s, Cell Signaling).

### Fluorescence Image Quantification

Fluorescence staining was quantified using ImageJ (NIH). Images were converted to 8-bit and analyzed using identical settings across all samples. A fixed-size region of interest (ROI) was drawn around each cell or area of interest, and integrated fluorescence intensity was measured. Background fluorescence from an unstained adjacent region was subtracted from each measurement. Fluorescence values were then normalized to the control group and expressed relative to control.

### Human Islet Proteomics and Pulldown Assays

For the studies on the differential protein response to harmine, human islets in **Fig. 4**, three different human islet preparations were treated for 72 hours with vehicle (0.1% DMSO) vs. harmine (10 μM), then lysed using lysis buffer containing 200 mM EPPS (pH 8.5) and 8 M urea, with complete protease and PhosSTOP phosphatase inhibitors, then flash-frozen then flash frozen and shipped to the Proteomics Facility at Harvard Medical School Center for Mass Spectrometry. There, samples were subjected to the facility’s standardized workflow. Peptides were purified and analyzed by liquid chromatography– tandem mass spectrometry (LC-MS/MS). Raw spectra were processed with the facility’s established pipeline, including peptide identification against the human UniProt database and label-free quantification. Protein-level abundances were generated after peptide-spectrum matching, false-discovery rate control (1% FDR at peptide and protein levels), and normalization using the Center’s standard procedures. The raw protein expression matrix was then log2-transformed and median-centered prior to analysis. The dataset included nine donor pairs, each consisting of three samples processed across different batches. Differential protein expression was assessed on 7,687 proteins detected in at least two of the three batches. LIMMA models were fitted using a design matrix including treatment condition (Harmine vs DMSO) and donor pair ID. Batch correction was not applied because each batch was perfectly confounded with donor identity, and including both would make the design matrix singular; the paired model fully accounts for between-pair differences. The Harmine–vs–DMSO contrast was estimated using empirical Bayes moderation, and the full DPE table was extracted with FDR correction.

For the pulldown studies (**Fig. 6A-E**), harmine was purchased from (Sigma, St. Louis). 2-2c and compound 15 were synthesized at Mount Sinai as described^26,27^. Pulldown of human islet lysates, magnetic bead coupling to compound 15 and graded elution, followed by shotgun proteomics were performed at Evotec International GmbH, Munich, Germany.

### Protein Purification For Thermal Shift Assay

A plasmid encoding human PRKAR1A plasmid was a gift from William Hahn & David Root (Addgene plasmid # 23741; http://n2t.net/addgene:23741; RRID:Addgene_23741). The plasmid used to express PRKACA in mammalian cells was used as a template for bacterial expression. We subcloned both the regulatory (PRKAR1A) and catalytic subunit (PRKACA) separately into His-Sumo N-terminal tagged vectors. For expression, the plasmids were introduced into LOBSTR cells, a modified BL21 (DE3) strain. For PRKACA, the plasmid was co-transformed with lambda phosphatase (in a pCDF vector) to reduce toxicity. Terrific Broth (TB) media supplemented with kanamycin for PRKAR1A and both kanamycin and streptomycin for PRKACA at 50 mg/ml final concentration was inoculated with 1:1000 dilution of an overnight culture. The culture was grown at 37°C and the temperature was reduced to 16°C once the OD_600_ reached 1.5. Protein production was induced overnight by adding IPTG to a final concentration of 0.4 mM at 16°C. The cells were harvested the next day at 4°C and then resuspended in TBS (20 mM Tris pH 8.0 and 250 mM NaCl, supplemented with 1 mM PMSF and 0.1 mg/ml Lysozyme). The cells were then lysed by sonication. The supernatant was supplemented with 40 mM imidazole and then purified by IMAC. After batch binding, the column was washed with TBS containing 50 mM imidazole and then eluted with TBS containing 250 mM Imidazole and 10% glycerol. After cleaving the His-Sumo tag with Sumo protease overnight, the protein was further purified through Superdex-200 gel filtration sizing in TBS containing 20 mM Tris pH 8.0 and 150 mM NaCl.

### Protein Thermal Shift Assay for PRKAR1A and PRKACA

The thermal shift assay to assess the harmine and cAMP binding to PRKAR1A and PRKACA, individually and as a complex was performed on a QuantStudio 3 real-time PCR system (Applied Biosystems). In the final reaction volume of 20 mL, proteins were diluted to a final concentration of 0.5 mg/ml, in TBS buffer. Both cAMP and Mg-ATP in water were diluted to a final 1 mM concentration. Harmine was added to a final concentration of 0.1 mM at 1% DMSO from 1 mM in 10 % DMSO stock. 10x SYPRO orange (Applied Biosystems) was prepared by adding 0.5 ml of 5000x to 250 mL of TBS buffer and in the final reaction mix, 2 ml of this dye was added. The reactions were kept at 25°C for 2 min and then heated from 25°C to 95°C with a rate of 0.05°C/S. The change in the fluorescence intensities of SYPRO orange was monitored as a function of the temperature and analyzed by Protein Thermal Shift Software 1.3. Each reaction was performed in triplicate.

### Statistics

Statistical analyses were performed using Student’s 2-tailed paired *t-*test as described in the Figure Legends. P values less than 0.05 were considered to be significant.

## Supplementary Figures

**Supplemental Fig. 1.**
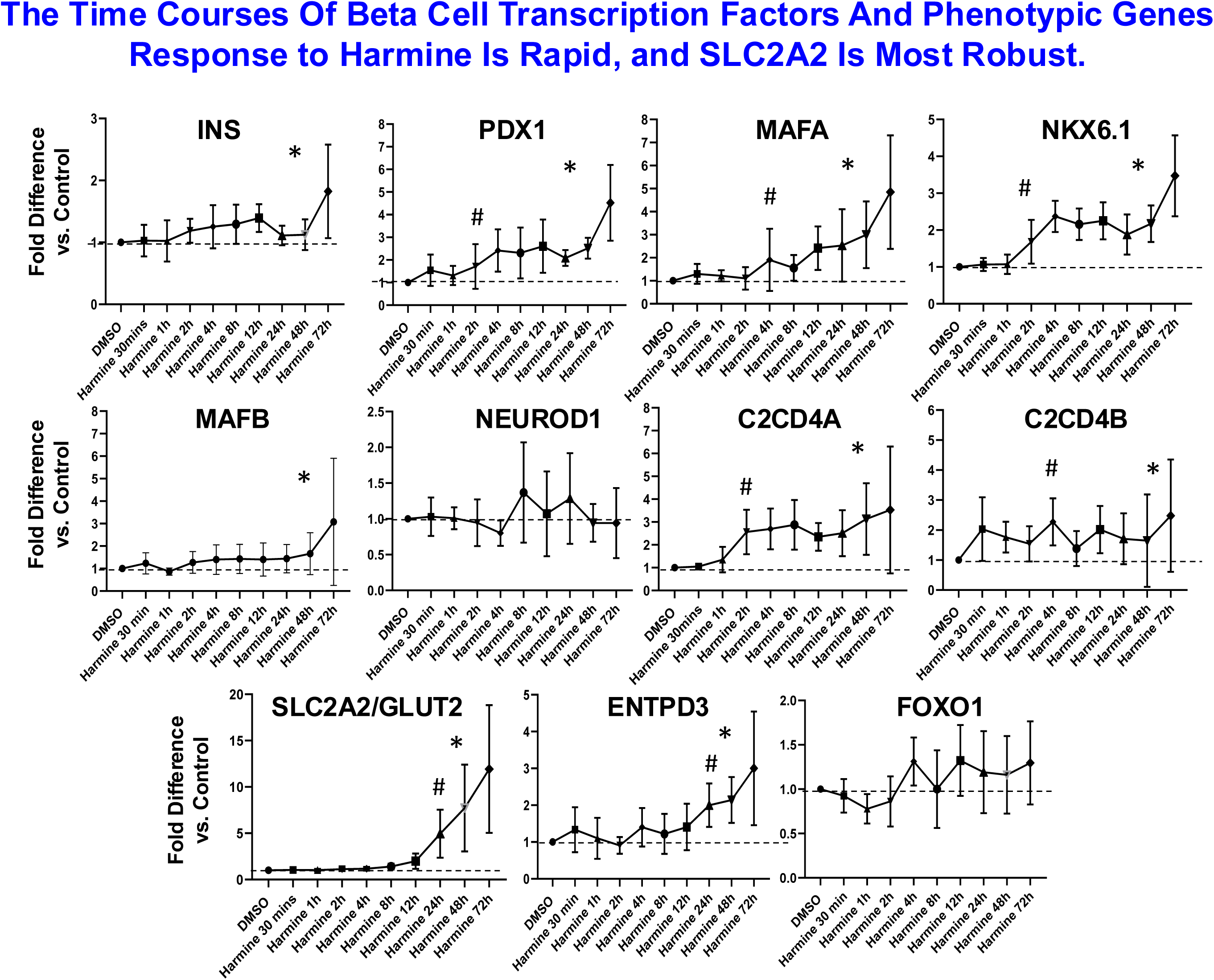
Time Course of Increase in Selected Differentiation Markers. Note that PDX1, MAFA, NKX6.1 and C2CD4A rise within 2-8 hours, whereas MAFB, ENTPD3 and SLC2A2 (GLUT2) require 12-24 hours to respond. Note also that other transcription factors (NEUROD1, FOXO1, C2CD4B) show little response. Note that Insulin mRNA changes only modestly, as reported previously, although insulin secretion increased in vitro and in vivo in response to harmine^17,18^. Asterisks represent P<0.05 for the complete dataset vs. time 0 by oneway ANOVA, and the # symbols indicate the earliest time point at which the factor in question becomes elevated, as determined by paired T-tests vs. time 0.

**Supplemental Fig 2.**
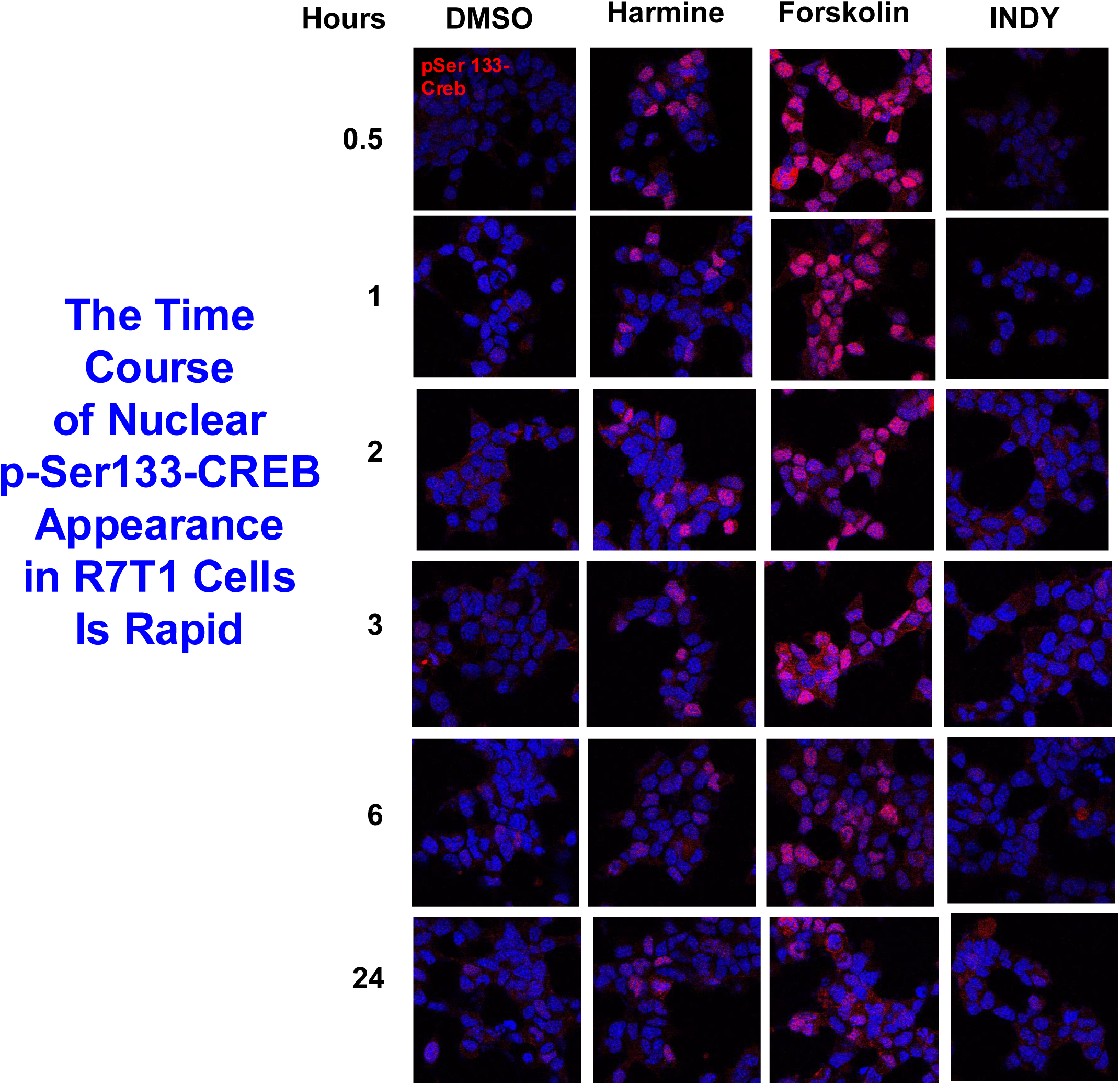
Time Course of Harmine and Forskolin Induction of ^Ser133^P-CREB In Mouse R7T1 Cells. The point is that harmine, like forskolin, induces ^Ser133^P-CREB phosphorylation in R7T1 cells, beginning within 30 min, and gradually declining over 2-3 hours, paralleling the effects of forskolin.

## Supplementary Tables

**Supplementary Table 1. Human Islet Organ Donor Summary.**

**Supplementary Table 2. Primers Used For PCR Experiments.**

**Supplementary Table 3. Human Islet Proteomic Data.**

